# Dynein-dynactin segregate meiotic chromosomes in *C. elegans* spermatocytes

**DOI:** 10.1101/2020.10.19.345041

**Authors:** Daniel J. Barbosa, Vanessa Teixeira, Joana Duro, Ana X. Carvalho, Reto Gassmann

## Abstract

The dynactin complex is an essential co-factor of the microtubule-based motor dynein. Dynein-dynactin have well-documented roles in spindle assembly and positioning during *C. elegans* female meiosis and embryonic mitosis, while dynein-dynactin’s contribution to male meiosis has not been investigated. Here, we characterize the G33S mutation in DNC-1’s N-terminal microtubule binding domain, which corresponds to G59S in the human dynactin subunit p150/Glued that causes motor neuron disease. In spermatocytes, *dnc-1(G33S)* delays spindle assembly and penetrantly inhibits anaphase spindle elongation in meiosis I, which prevents homologous chromosome segregation and generates aneuploid sperm with an extra centrosome. Consequently, embryos produced by *dnc-1(G33S)* hermaphrodites exhibit a high incidence of tetrapolar mitotic spindles, yet *dnc-1(G33S)* embryos with bipolar spindles proceed through early mitotic divisions without errors in chromosome segregation. Deletion of the DNC-1 N-terminus shows that defective meiosis in *dnc-1(G33S)* spermatocytes is not due to DNC-1’s inability to interact with microtubules. Rather, our results suggest that the DNC-1(G33S) protein, which is aggregation-prone *in vitro*, is less stable in spermatocytes than the early embryo, resulting in different phenotypic severity in the two dividing tissues. Thus, the unusual hypomorphic nature of the *dnc-1(G33S)* mutant reveals that dynein-dynactin drive meiotic chromosome segregation in spermatocytes and illustrates that the extent to which protein misfolding leads to loss of function can vary significantly between cell types.

## INTRODUCTION

Eukaryotic chromosome segregation during cell division is accomplished by a bipolar microtubule-based spindle that connects to chromosomes via their kinetochores. Chromosomes can be segregated by moving towards spindle poles (anaphase A) and by virtue of the two spindle poles moving apart (spindle elongation, anaphase B). Studies in diverse model organisms show that most cells use a combination of anaphase A and B (Maiato and Lince-Faria, 2010), while some cells, such as the one-cell embryo of the nematode *Caenorhabditis elegans*, rely primarily on spindle elongation to segregate chromosomes. Two types of forces have been shown to drive spindle elongation (Scholey et al., 2016): microtubules that are present between segregating chromosomes (referred to as interpolar, interchromosomal, central spindle, or midzone microtubules) can generate outward-directed pushing forces through a combination of filament sliding and changes in microtubule dynamics, and, in spindles with centrosomes, interactions between the cell cortex and astral microtubules can generate forces that pull on spindle poles. The latter mechanism has been shown to require cortical localization of the motor complex cytoplasmic dynein 1 (dynein), whose motility is directed towards the microtubule minus ends anchored at centrosomes. Dynein acts at various other subcellular locations besides the cortex, such as the nuclear envelope, kinetochores, and spindle poles, and is involved in multiple aspects of spindle assembly, including the separation of the nucleus-associated centrosomes in prophase and the establishment of kinetochore-microtubule interactions in prometaphase (Raaijmakers et al., 2014). Dynein-dependent pulling forces at the cortex are universally used in eukaryotes to position and orient the spindle (Kiyomitsu, 2019). By contrast, the contribution of cortical pulling forces to anaphase varies significantly between species, and it can also vary between different cells in the same organism, as exemplified by *C. elegans:* in the one-cell embryo, dynein-dependent pulling forces at the cortex segregate chromosomes by elongating the spindle, while midzone-derived pushing forces are responsible for acentrosomal chromosome segregation during female meiosis (Laband et al., 2017; McNally et al., 2016). During *C. elegans* male meiosis, which occurs in the presence of centrosomes, chromosome segregation is achieved primarily by spindle elongation (Shakes et al., 2009; Fabig et al., 2020), but dynein’s role in spermatocytes remains uncharacterized.

Both dynein localization and dynein activity critically depend on its co-factor dynactin. The dynactin complex is built around a short Arp1-based actin filament, whose interaction with the dynein heavy chain (HC) dimer results in allosteric activation of processive motility (Carter et al., 2016). The interaction between dynein and dynactin is inherently weak and requires stabilization by adaptor proteins that recruit dynein-dynactin to their diverse sites of action such as the cell cortex. The largest dynactin subunit, p150/Glued, is part of the shoulder complex, which is situated on one side of the Arp1 filament near its barbed end (Fig. 1A). The N-terminal region of p150/Glued forms a long projecting arm that features two coiled-coil regions (CC1 and CC2) and a highly conserved N-terminal cytoskeleton-associated protein glycine-rich (CAP-Gly) domain that binds microtubules and the microtubule plus-end tracking proteins (+TIPs) CLIP-170 and end-binding (EB) protein. A patch of basic residues adjacent to the CAP-Gly domain also binds microtubules, while CC1 binds dynein intermediate chain (IC). While p150/Glued is an essential dynactin subunit, its CAP-Gly domain has more specialized functions, namely in ensuring robust initiation of dynein motility (Moore et al., 2009; Moughamian et al., 2012; Lloyd et al., 2012; McKenney et al., 2016; Barbosa et al., 2017).

**Figure 1.**
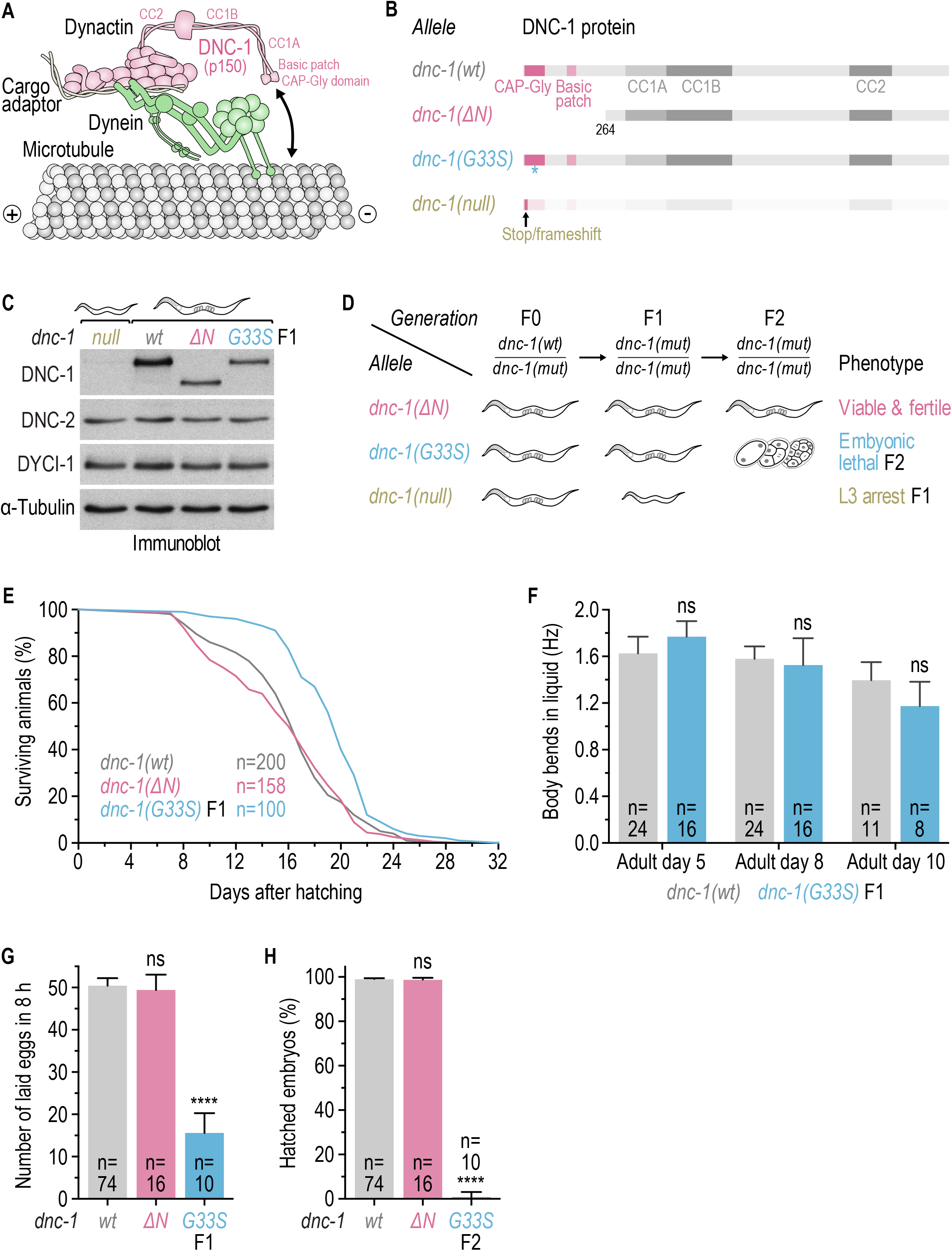
The penetrant lethality of *dnc-1(G33S)* F2 progeny contrasts with the normal viability of an N-terminal *dnc-1* deletion. **(A)** Cartoon of the dynein-dynactin transport machine on a microtubule. The DNC-1 subunit (p150/Glued) extends from the dynactin filament and binds the microtubule through its N-terminal CAP-Gly domain and an adjacent basic patch. **(B)** Schematic of DNC-1 proteins expressed by the mutants examined in this study. Opaqueness indicates that no protein is produced in the *dnc-1(null)* mutant. **(C)** Immunoblot of hermaphrodites with antibodies against DNC-1, DNC-2 (p50/dynamitin), DYCI-1 (dynein intermediate chain), and α-tubulin, which serves as the loading control. **(D)** Fate of homozygous *dnc-1* mutant progeny (F1 and F2) starting from heterozygous mothers (F0). **(E)** Life span curves with *n* denoting the number of hermaphrodites examined. **(F)** Locomotion of adult hermaphrodites assessed by determining body bending frequency in liquid medium (mean ± CI 95%) with *n* denoting the number of animals examined. Statistical significance (wild-type versus mutant *dnc-1)* was determined by ANOVA on ranks (Kruskal-Wallis nonparametric test) followed by Dunn’s multiple comparison test. ns = not significant, *P* > 0.05. **(G)** Number of eggs laid by adult hermaphrodites in an 8h window (mean ± CI 95%) with *n* denoting the number of mothers whose progeny was counted. Statistical significance (wild-type versus mutant *dnc-1)* was determined by ANOVA on ranks (Kruskal-Wallis nonparametric test) followed by Dunn’s multiple comparison test. \*\*\*\**P* < 0.0001; ns = not significant, *P* > 0.05. **(H)** Embryonic viability (mean ± CI 95%) with *n* denoting the number of hermaphrodite mothers whose progeny was examined. Statistical significance (wild-type versus mutant *dnc-1)* was determined by ANOVA on ranks (Kruskal-Wallis nonparametric test) followed by Dunn’s multiple comparison test. \*\*\*\**P* < 0.0001; ns = not significant, *P* > 0.05.

Point mutations in the CAP-Gly domain of p150/Glued cause Perry syndrome and hereditary motor neuropathy 7B (HMN7B; also known as distal spinal and bulbar muscular atrophy), adult-onset autosomal dominant neurodegenerative diseases that primarily affect dopaminergic neurons in the substantia nigra and motor neurons, respectively (Puls et al., 2003; Farrer et al., 2009). The disease mutations reduce p150/Glued’s affinity for microtubules and +TIPs, at least in part by disrupting the CAP-Gly domain fold (Puls et al., 2003; Levy et al., 2006; Farrer et al., 2009; Ahmed et al., 2010; Moughamian et al., 2012; Lloyd et al., 2012). The HMN7B mutation G59S is predicted to result in particularly severe CAP-Gly domain misfolding, because, unlike the surface-exposed residues mutated in Perry Syndrome (F52L, K56R, G67D, G71A, G71E, G71R, T72P, Q74P, Y78C), G59 is located in the domain’s hydrophobic core. p150/Glued containing the HMN7B mutation is prone to aggregation when overexpressed in cells and when synthesized *in vitro* (Levy et al., 2006; Lloyd et al., 2012; Moughamian et al., 2012), and protein levels of mutated endogenous p150/Glued are reduced in the mouse (G59S), *D. melanogaster* (G38S), and *C. elegans* (G33S), indicating that the mutation affects protein stability (Lai et al., 2007; Lloyd et al., 2012; Barbosa et al., 2017). While p150/Glued(G59S) is readily incorporated into the dynactin complex (Lai et al., 2007; Levy et al., 2006; Chevalier-Larsen et al., 2008), the mutation was reported to reduce the association between dynactin and dynein in pull-down experiments from cell lysate (Moughamian et al., 2012). Thus, while both Perry Syndrome and HMN7B mutations impair CAP-Gly domain function, the HMN7B mutation has additional adverse consequences for dynactin function that are related to its destabilizing effect on the CAP-Gly domain fold. This is likely to be relevant for understanding why mutations of residues that are in close proximity cause distinct neurodegenerative diseases.

We previously found that the Perry syndrome mutations F26L (human F52L) and G45R (human G71R) in the *C. elegans* p150/Glued ortholog DNC-1 mildly perturb dynein-dynactin function in the one-cell embryo (Barbosa et al., 2017). Specifically, *dnc-1(F26L)* and *dnc-1(G45R)* embryos exhibit delays in the centration and rotation of the male pronucleus, which is associated with the female pronucleus and the two centrosomes. Here, we characterize the HMN7B mutation G33S and show that it behaves as an unusual hypomorph whose phenotypic severity varies significantly between the early embryo and spermatocytes. In the zygote and during early embryogenesis, *dnc-1(G33S)* produces mild defects in spindle positioning during mitotic divisions that resemble the defects of Perry Syndrome mutants and are also observed with an N-terminal deletion mutant, *dnc-1(ΔN)*, which lacks the CAP-Gly domain and the basic patch. By contrast, chromosome segregation in spermatocytes is specifically and severely affected in *dnc-1(G33S)* animals. The *dnc-1(G33S)* mutant reveals that, unlike chromosome segregation during female meiosis, chromosome segregation during male meiosis requires dynein-dynactin activity in anaphase. Specifically, our results suggest that spindle elongation during male meiosis is accomplished exclusively by dynein-mediated cortical pulling forces, in agreement with recent ultrastructural analysis showing that anaphase spindles lack a typical midzone architecture that could generate pushing forces (Fabig et al., 2020). Finally, we provide evidence that tissue-specific differences in DNC-1(G33S) stability may be the underlying cause for the phenotypic differences between the *dnc-1(G33S)* and *dnc-1(ΔN)* mutant.

## RESULTS

### Dynactin DNC-1’s N-terminal microtubule binding region is dispensable for viability

We previously found that *C. elegans* strains carrying the Perry syndrome mutations F26L and G45R (F52L and G71R in human p150/Glued, respectively) in DNC-1’s CAP-Gly domain are homozygous viable, even in the context of a DNC-1 splice isoform that lacks the basic patch (Barbosa et al., 2017). This raised the possibility that microtubule binding by dynactin may be dispensable for viability, but we could not rule out that the mutant CAP-Gly domain retained residual affinity for microtubules. We therefore used genome editing to generate a deletion mutant, *dnc-1(ΔN)*, which expresses a truncated version of DNC-1 that starts at methionine 264 (Fig. 1A, B). The deleted region includes the CAP-Gly domain (residues 5-69), the adjacent basic patch (residues 140-169), and part of the linker (residues 170-325) that precedes the first coiled-coil region, CC1 (residues 326-664). *dnc-1(ΔN)* hermaphrodites could be propagated in a homozygous state, showed no significant decrease in embryonic viability, had a normal life span (wild type N2 average 16.4 ± 0.6 days, *dnc-1(ΔN)* 15.8 ± 0.8 days, *p* = 0.9096), and were fully fertile (Fig. 1B, D, E, G, H). These results demonstrate that the microtubule binding activity of *C. elegans* dynactin is dispensable for viability and fertility.

### The DNC-1 G33S mutation results in penetrant embryonic lethality

We next compared the *dnc-1(ΔN)* mutant with the CAP-Gly domain mutant G33S, which corresponds to the HMN7B mutation G59S in human p150/Glued (Fig. 1B). Immunoblotting of homozygous hermaphrodites showed that DNC-1(ΔN) and DNC-1(G33S) are expressed at comparable levels, which are lower than in wild-type N2 hermaphrodites (Fig. 1C). Reverse transcription PCR analysis further showed that *dnc-1* mRNA levels are unaffected in either mutant and that the *dnc-1(G33S)* mutation does not interfere with the generation of N-terminal splice variants (Fig. S1A-D). Although *dnc-1(G33S)* and *dnc-1(ΔN)* hermaphrodites express the same amount of mutant protein, the *dnc-1(G33S)* mutation cannot not be propagated when homozygous (Fig. 1D). We therefore maintained the *dnc-1(G33S)* mutation in a heterozygous state using a GFP-marked balancer. Hermaphrodites homozygous for *dnc-1(G33S)*, identified by the absence of GFP expression in the pharynx, grew to adulthood and produced embryos, albeit at a reduced rate, and these embryos failed to hatch (Fig. 1D, G, H). This phenotype is distinct from that of a *dnc-1* null mutant, which invariably results in developmental arrest of homozygous hermaphrodites at the larval L3 stage (Fig. 1B-D). Despite producing inviable embryos, homozygous *dnc-1(G33S)* hermaphrodites lived slightly longer than wild-type N2 hermaphrodites (19.6 ± 0.7 days, *p* < 0.0001) and performed similar to wild-type N2 in a liquid locomotion assay during the course of aging (Fig. 1E, F). These results show that *dnc-1(G33S)* is not a null mutant and that the DNC-1(G33S) protein retains sufficient functionality to support development and partial fertility.

### An extra paternally derived centrosome causes tetrapolar mitotic spindles in one-cell *dnc-1(G33S)* embryos

Given that *dnc-1(ΔN)* animals are fully viable, the penetrant embryonic lethality caused by the G33S mutation in the F2 generation cannot be explained by the lack of CAP-Gly domain function. To understand why the two mutants behave differently, we monitored the first embryonic division by live imaging using mCherry::HIS-11 (histone H2B) and GFP::TBB-2 (β-tubulin) to mark chromosomes and microtubules, respectively. After centration and rotation of the nucleus-centrosome complex (NCC), control embryos assembled a bipolar spindle that was correctly orientated along the anterior-posterior axis (Fig. 2A). In *dnc-1(ΔN)* embryos, a mis-oriented bipolar spindle assembled in the posterior half as a consequence of delayed NCC centration/rotation. The delay was slightly more pronounced than that previously observed in *dnc-1(G45R)* and *dnc-1(F26L)* embryos (Barbosa et al., 2017), consistent with the decreased DNC-1(ΔN) levels detected by immunoblot. Nevertheless, spindle orientation eventually corrected during prometaphase and chromosomes segregated in anaphase without apparent errors. By contrast, depletion of DNC-1 by RNA interference (RNAi) inhibited centrosome separation in prophase, which resulted in monopolar spindle formation in prometaphase and thus prevented chromosome segregation, consistent with prior work (Gönczy et al., 1999; Gama et al., 2017). Embryos produced by hermaphrodites homozygous for *dnc-1(G33S)*, i.e. F2 *dnc-1(G33S)* embryos, could be grouped into two phenotypic classes (Fig. 2A, B). In 18 out of 42 embryos (43 %), the phenotype resembled that of *dnc-1(ΔN)* embryos: a bipolar spindle assembled after delayed NCC centration/rotation and chromosomes segregated correctly in anaphase. The remaining 24 embryos (57 %) formed a tetrapolar spindle, which predictably caused severe chromosome mis-segregation. Extra spindle poles were never observed in *dnc-1(ΔN)* embryos (n = 20) or in embryos produced by hermaphrodites heterozygous for *dnc-1(G33S)* (n = 18). In the *dnc-1(G45R)* mutant, only 6 out of 45 embryos (13 %) formed a tetrapolar spindle (Fig. 2B).

**Figure 2.**
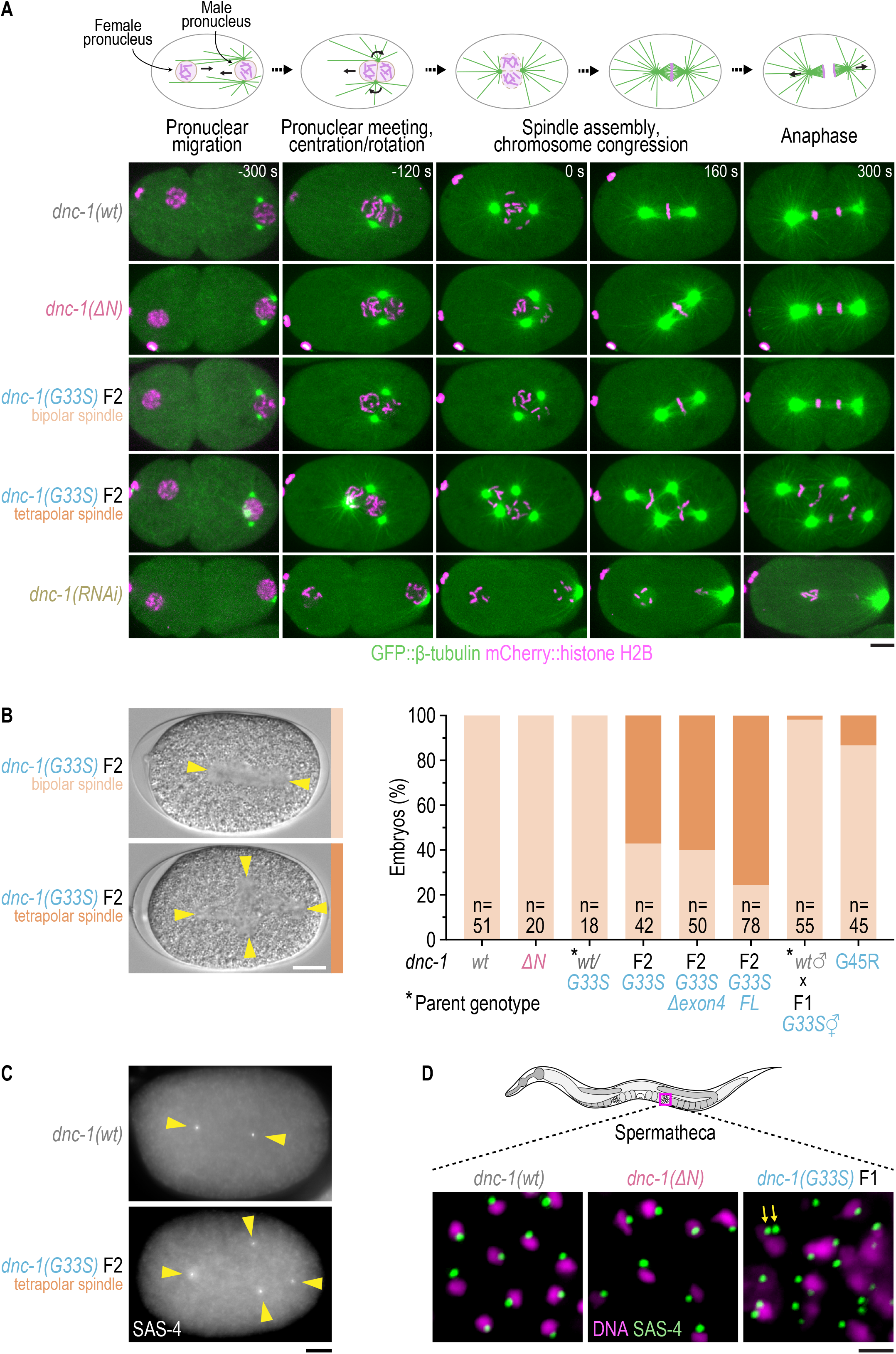
An extra paternally derived centrosome generates tetrapolar spindles in one-cell *dnc-1(G33S)* embryos. **(A)** Stills from time-lapse fluorescence imaging of the first embryonic division from pronuclear migration until anaphase. Time point 0 s corresponds to nuclear envelope breakdown. *F2* denotes the second generation of homozygous *dnc-1(G33S)* progeny, as described in Figure 1D. Scale bar, 5 μm. **(B)** (*left*) Stills from time-lapse differential interference contrast (DIC) imaging of the first embryonic division. Arrowheads mark the spindle poles. Scale bar, 10 μm. (*right*) Frequency of bipolar versus tetrapolar spindles in one-cell embryos, determined by time-lapse DIC imaging, with *n* denoting the number of embryos examined. **(C)** Immunofluorescence images of one-cell embryos stained with an antibody against the centriole component SAS-4. Arrowheads mark the centrioles. Scale bar, 5 μm. **(D)** Immunofluorescence images of sperm in adult hermaphrodites stained with an antibody against the centriole component SAS-4. DNA was visualized with DAPI. Arrows point to two centrosomes in a *dnc-1(G33S)* mutant sperm cell. Scale bar, 2 μm.

We previously showed that DNC-1’s basic patch is subject to alternative splicing (Barbosa et al., 2017). To determine how DNC-1 splice isoforms influence the tetrapolar spindle phenotype caused by the G33S mutation, we generated strains in which the basic patch is either always present (full-length isoform, *FL*) or always absent (*Δexon4* isoform) (Fig. S1E). 76 % of *dnc-1(G33S) FL* embryos (n = 78) and 60 % of *dnc-1(G33S) Δexon4* embryos (n = 50) assembled a tetrapolar spindle (Fig. 2B). Thus, the tetrapolar spindle defect is enhanced when DNC-1(G33S) is forced to retain its basic patch, while removal of the basic patch has no effect.

Immunofluorescence analysis with an antibody against the centriole component SAS-4 confirmed that each of the four spindle poles in *dnc-1(G33S)* embryos contained a centrosome (Fig. 2C). In sperm of homozygous *dnc-1(G33S)* hermaphrodites, the SAS-4 antibody detected either one or two centrosomes, while wild-type and *dnc-1(ΔN)* sperm never contained more than one centrosome (Fig. 2D). The single centrosome donated to the zygote by wild-type sperm consists of two centrioles that separate to form the two spindle poles in the first embryonic division. This suggests that tetrapolar spindles in *dnc-1(G33S)* embryos arise because of an extra paternally derived centrosome. Consistent with this, the tetrapolar spindle defect could be rescued by mating homozygous *dnc-1(G33S)* hermaphrodites with wild-type males (Fig. 2B). Quantitative analysis of pronuclear migration revealed that the NCC centration defect was slightly less pronounced in *dnc-1(G33S)* embryos than in *dnc-1(ΔN)* embryos, irrespective of whether *dnc-1(G33S)* embryos contained two or four centrosomes (Fig. 3A-E). In contrast to the tetrapolar spindle defect, wild-type sperm failed to rescue the NCC centration defect in *dnc-1(G33S)* embryos (Fig. 3D).

**Figure 3.**
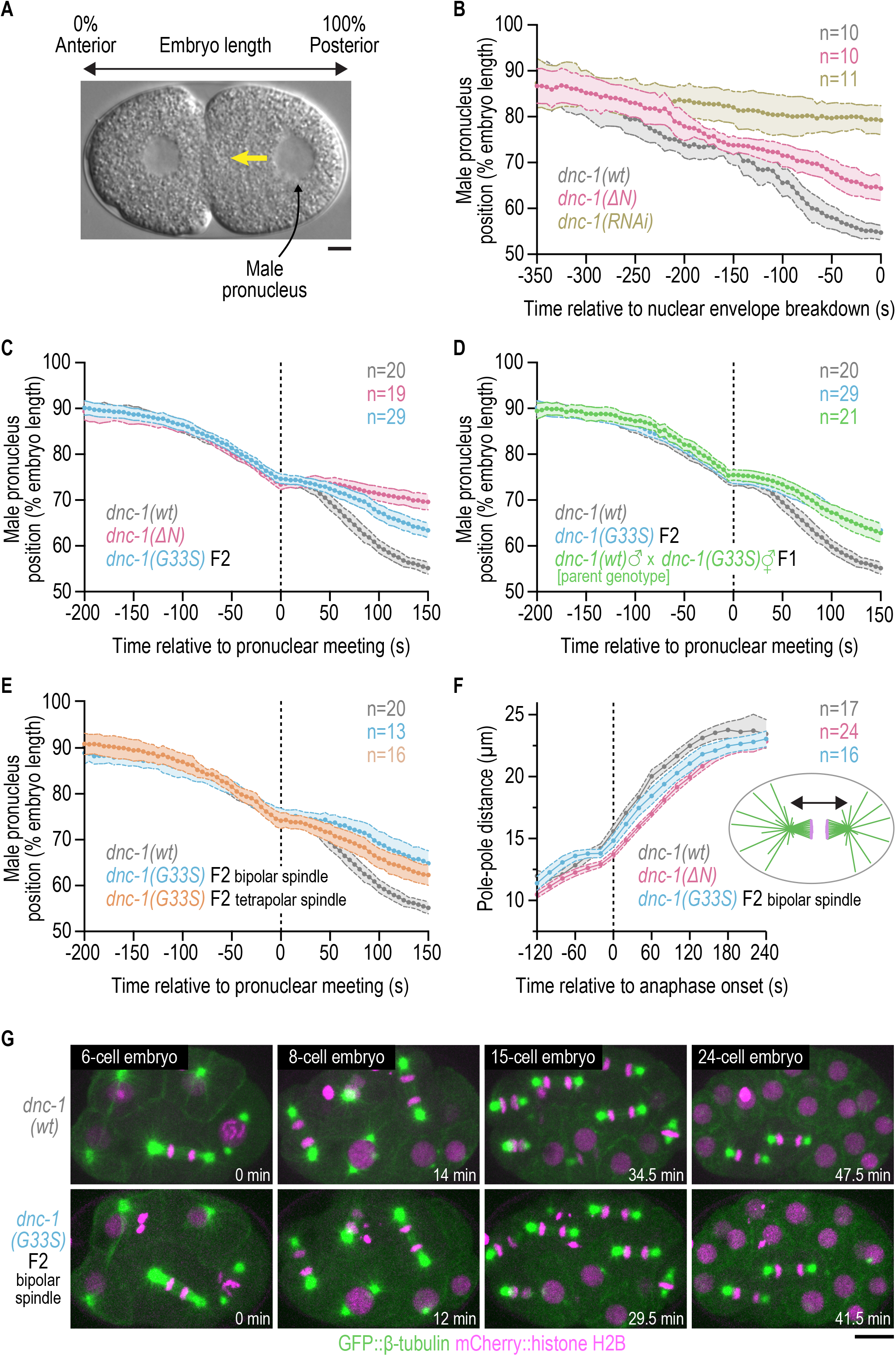
One-cell *dnc-1(G33S)* embryos with bipolar spindles exhibit milder defects than *dnc-1(ΔN)* embryos and complete early mitotic divisions without chromosome segregation errors. **(A)** Still image from time-lapse DIC imaging in a one-cell embryo. The yellow arrow indicates the migration of the male pronucleus along the anterior-posterior axis, which is plotted in panels (*B) - (E)*. Scale bar, 5 μm. **(B) - (E)** Position of the male pronucleus (mean ± 95 CI) versus time with *n* denoting the number of embryos examined. **(F)** Spindle length (mean ± 95 CI) versus time in one-cell embryos with *n* denoting the number of embryos examined. **(G)** Still images from time-lapse fluorescence imaging of early embryonic divisions. Time point 0 min corresponds to mid-anaphase in the EMS cell. Scale bar, 10 μm.

We conclude that *dnc-1(G33S)* embryos experience two defects of distinct origin: a paternally derived defect, characterized by the presence of an extra centrosome that results in tetrapolar spindle formation, and a maternally derived defect, characterized by delayed NCC centration/rotation. Because only the latter defect is present in *dnc-1(ΔN)* embryos, the paternally derived defect in *dnc-1(G33S)* embryos cannot be ascribed to DNC-1(G33S)’s inability to bind microtubules.

### The DNC-1 G33S mutation inhibits meiotic chromosome segregation in spermatocytes

To understand why the sperm of homozygous *dnc-1(G33S)* mutants contains an extra centrosome, we monitored spermatogenesis in L4 hermaphrodites by live-cell imaging *in situ* using the mCherry::histone H2B and GFP::β-tubulin probes (Fig. 4A). As spermatocytes approached late prophase of meiosis I, microtubule-nucleating centrosomes could be clearly distinguished using the GFP::β-tubulin signal. Just like control and *dnc-1(ΔN)* spermatocytes, *dnc-1(G33S)* spermatocytes invariably contained two centrosomes (Fig. 4B), indicating that the mitotic divisions of syncytial stem cell nuclei at the distal end of the gonad had occurred without major defects. At the time of nuclear envelope breakdown (NEB), centrosomes were positioned on opposite sides of the nucleus in control and *dnc-1(ΔN)* spermatocytes, while centrosomes in *dnc-1(G33S)* spermatocytes were frequently incompletely separated at NEB.

**Figure 4.**
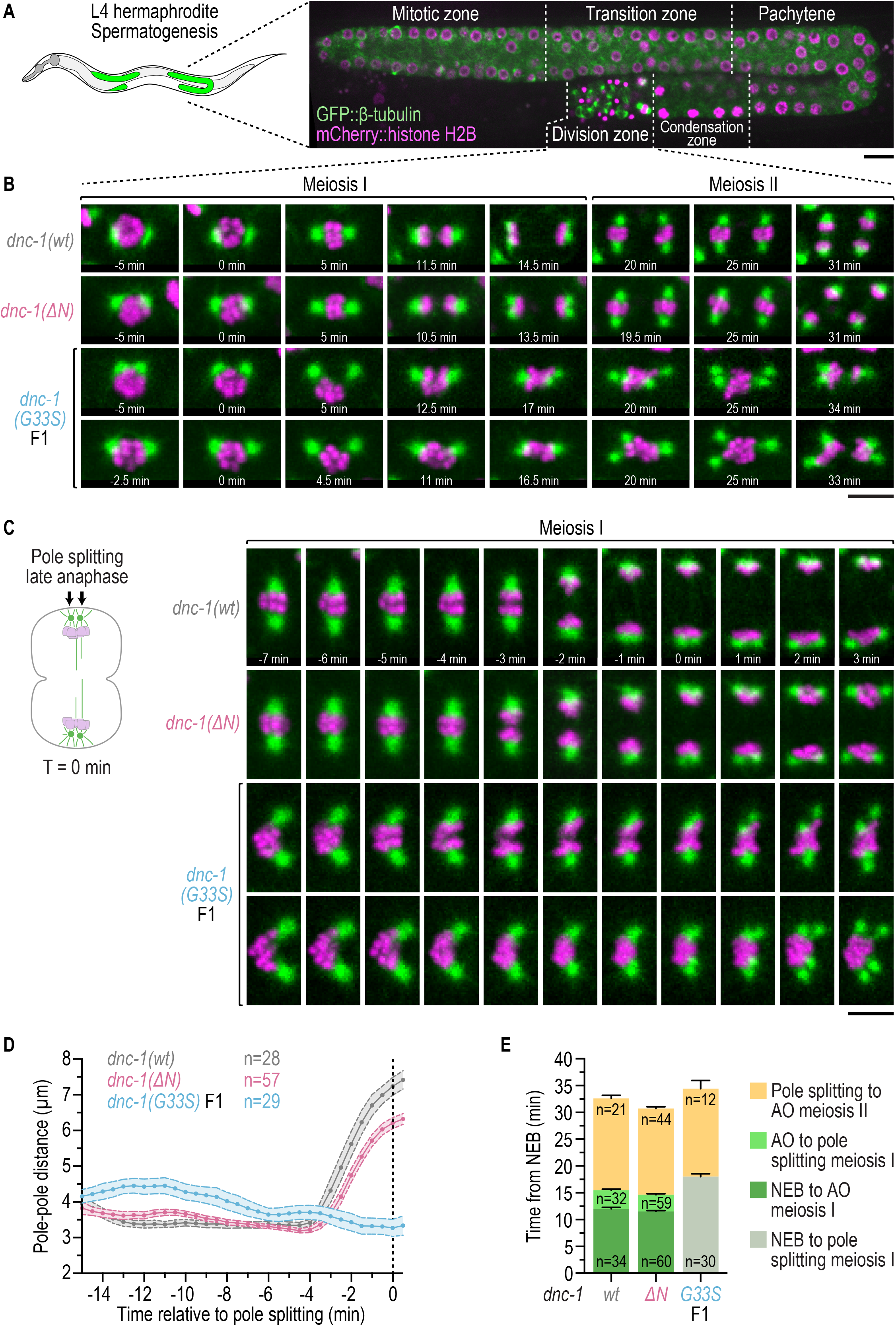
The DNC-1(G33S) mutation inhibits meiotic chromosome segregation during spermatogenesis in hermaphrodites. **(A)** Fluorescence image of the sperm-producing gonad in an L4 hermaphrodite. The division zone is the focus of panels *(B) - (E)*. Scale bar, 10 μm. **(B)** Stills from time-lapse fluorescence imaging of meiosis I and II, starting at the end of prophase. Time point 0 min corresponds to the first discernable chromosome movements following nuclear envelope breakdown. Two examples are shown for *dnc-1(G33S)*. Scale bar, 5 μm. **(C)** Stills from time-lapse fluorescence imaging of chromosome segregation during meiosis I. Time point 0 min corresponds to the onset of pole splitting in late anaphase I, as depicted in the cartoon. Two examples are shown for *dnc-1(G33S)*. Scale bar, 5 μm. **(D)** Spindle length (mean ± 95 % CI) versus time during meiosis I with *n* denoting the number of divisions examined. **(E)** Duration of meiosis intervals determined from time-lapse fluorescence imaging with *n* denoting the number of intervals scored. NEB, nuclear envelope breakdown (first discernable chromosome movements). AO, anaphase onset (first discernable separation of sister chromatids). Note that AO could not be reliably scored in the *dnc-1(G33S)* mutant.

Control and *dnc-1(ΔN)* spermatocytes reached metaphase I within 5 min after NEB, and anaphase I initiated 12.0 ± 0.3 min after NEB (Fig. 4B). By contrast, chromosomes struggled to congress in *dnc-1(G33S)* spermatocytes. Chromosomes typically moved away from partially separated centrosomes and became positioned on one side of the nascent spindle. The anti-poleward chromosome motion is likely a consequence of defects in establishing timely kinetochore-microtubule attachments that usually oppose polar ejection forces. Despite these initial problems, most *dnc-1(G33S)* spermatocytes eventually succeeded in incorporating chromosomes into a bipolar spindle whose length was similar to that measured in control and *dnc-1(ΔN)* spermatocytes (Fig. 4B-D). In control spermatocytes, the segregation of homologous chromosomes occurred primarily by anaphase B-type movement, during which the spindle elongated from 3.6 ± 0.1 μm to 7.2 ± 0.3 μm over a period of approximately 4 min (Fig. 4C, D). In *dnc-1(ΔN)* spermatocytes, anaphase I occurred successfully, albeit with slightly attenuated kinetics. By contrast, anaphase I completely failed in *dnc-1(G33S)* spermatocytes. Although homologous chromosomes could typically be observed to separate from each other, indicating timely loss of cohesion, there was no discernable spindle elongation; consequently, chromosomes remained unsegregated (Fig. 4B-D). In control and *dnc-1(ΔN)* spermatocytes, each spindle pole split into two during late anaphase of meiosis I in preparation for bipolar spindle assembly in meiosis II. Spindle pole splitting was only slightly delayed in the *dnc-1(G33S)* mutant (Fig. 4E), revealing that the absence of anaphase I is not an indirect consequence of cell cycle arrest. The failure of the first meiotic division in *dnc-1(G33S)* spermatocytes contrasts with the error-free mitotic divisions observed in their embryo progeny with bipolar spindles, where spindle elongation at the one-cell stage is only minimally affected and no defects in spindle assembly or chromosome segregation are evident in subsequent rounds of cell division (Fig. 3F, G).

Because *dnc-1(G33S)* spermatocytes failed to divide in meiosis I, tetrapolar spindles formed in meiosis II (Fig. 4B). Spindle poles initially moved away from the centrally positioned chromosomes towards the cell periphery, presumably because of polar ejection forces. As a consequence, spindle poles became separated from each other to a variable extent without forming proper bipolar spindles. In anaphase II, which initiated with similar timing as in control and *dnc-1(ΔN)* spermatocytes (Fig. 4E), chromosomes were randomly pulled towards spindle poles (Fig. 4B). Depending on the geometry of the aberrant anaphase II spindles, one or both centrosomes were subsequently incorporated into a budding spermatid (Fig. S2). Of note, even spermatids that correctly inherit one centrosome (resulting in bipolar spindle formation in the one-cell embryo) are likely to be aneuploid, which explains the penetrant embryonic lethality in *dnc-1(G33S)* progeny.

To complement our analysis of spermatogenesis in hermaphrodites, we also monitored spermatogenesis in young adult males using the same live-cell imaging approach. This analysis revealed that the defects caused by the *dnc-1(G33S)* mutation in males are essentially identical to those observed in hermaphrodites (Fig. 5A-F). We conclude that dynactin, and, by extension, dynein, is essential during anaphase I to drive chromosome segregation in spermatocytes.

**Figure 5.**
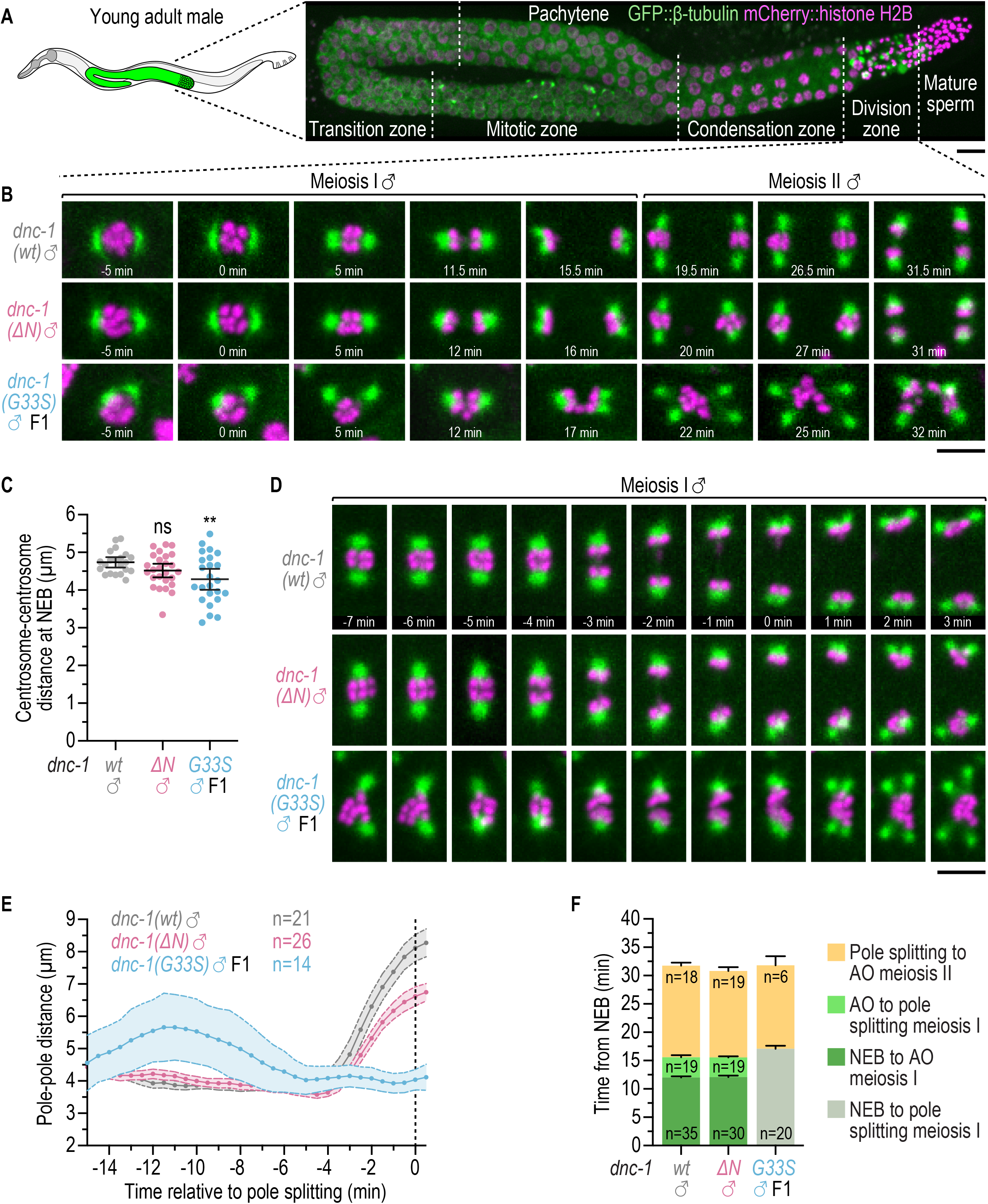
The DNC-1(G33S) mutation inhibits meiotic chromosome segregation in males. **(A)** Fluorescence image of the gonad in a young adult male. The division zone is the focus of panels *(B) - (F)*. Scale bar, 10 μm. **(B)** Stills from time-lapse fluorescence imaging of meiosis I and II, starting at the end of prophase. Time point 0 min corresponds to the first discernable chromosome movements following nuclear envelope breakdown. Scale bar, 5 μm. **(C)** Centrosome-centrosome distance (mean ± 95 % CI) measured just prior to nuclear envelope breakdown of meiosis I. Statistical significance (wild-type versus mutant *dnc-1)* was determined by one-way ANOVA followed by Dunnett’s multiple comparison test. \*\**P* < 0.01; ns = not significant, *P* > 0.05. **(D)** Stills from time-lapse fluorescence imaging of chromosome segregation during meiosis I. Scale bar, 5 μm. **(E)** Spindle length (mean ± 95 % CI) versus time during meiosis I with *n* denoting the number of divisions examined. **(F)** Duration of meiosis intervals as determined from time-lapse fluorescence imaging with *n* denoting the number of intervals scored. NEB, nuclear envelope breakdown (first discernable chromosome movement). AO, anaphase onset (first discernable separation of sister chromatids). Note that AO could not be reliably scored in the *dnc-1(G33S)* mutant.

### The DNC-1 G33S mutation de-localizes dynein in dividing spermatocytes but not in the early embryo

To gain insight into why the DNC-1(G33S) mutation causes a severe dynein loss-of-function phenotype in dividing spermatocytes but not in the dividing early embryo, we examined the localization of dynein using dynein heavy chain (DHC-1) endogenously tagged with GFP. In control embryos, GFP::DHC-1 was enriched at the nuclear envelope, the mitotic spindle, kinetochores, and the cell cortex, and a similar localization pattern was observed in control spermatocytes of young adult males (Fig. 6A-C; Fig. S3A-G). In *dnc-1(G33S)* embryos, GFP::DHC-1 was normally localized to the nuclear envelope, kinetochores, and the cell cortex, and exhibited slightly reduced signal on the spindle (Fig. S3A-G). By contrast, the GFP::DHC-1 signal was mostly diffuse in *dnc-1(G33S)* spermatocytes undergoing meiotic divisions, while dynein localization was unperturbed in *dnc-1(ΔN)* spermatocytes (Fig. 6A-C). We conclude that the severity of meiotic division defects in *dnc-1(G33S)* spermatocytes correlates with impaired dynein localization.

**Figure 6.**
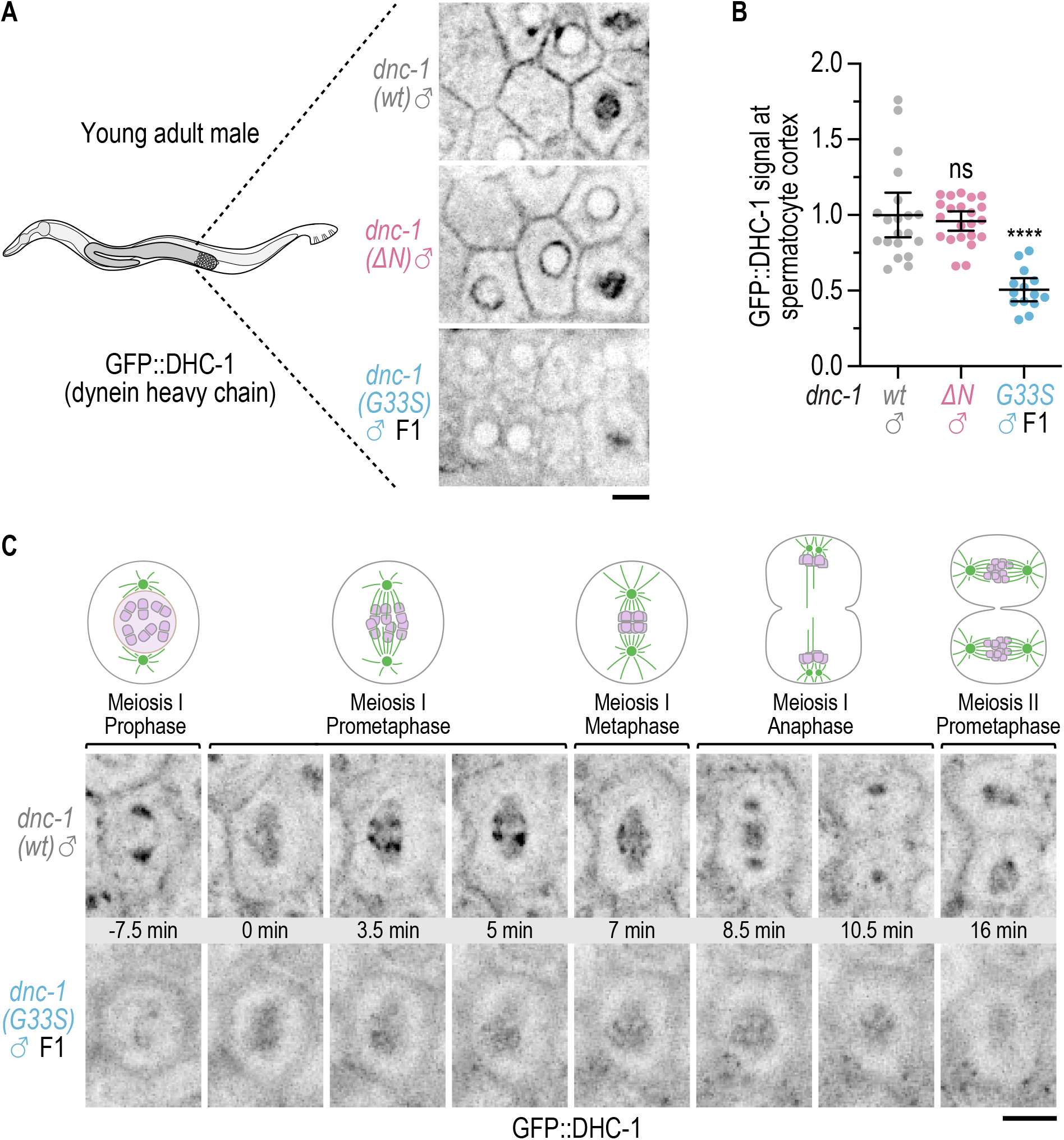
The DNC-1(G33S) mutation de-localizes dynein in dividing spermatocytes. **(A)** Fluorescence image of GFP::DHC-1 (dynein heavy chain) in the meiotic division zone of young adult males. Scale bar, 5 μm. **(B)** GFP::DHC-1 signal (mean ± 95 % CI) measured at the cortex of prophase spermatocytes and normalized to the wild-type control (*wt*). Statistical significance (wild-type versus mutant *dnc-1)* was determined by ANOVA on ranks (Kruskal-Wallis nonparametric test) followed by Dunn’s multiple comparison test. \*\*\*\**P* < 0.0001; ns = not significant, *P* > 0.05. **(C)** Stills from time-lapse fluorescence imaging of GFP::DHC-1 in meiosis I and II. Time point 0 min corresponds to the first image with discernable GFP signal in the spindle region following nuclear envelope breakdown. Scale bar, 5 μm.

### The G33S mutation renders the DNC-1 protein aggregation-prone *in vitro*

The G59S mutation in human p150/Glued results in mis-folding of the CAP-Gly domain, which makes the protein prone to aggregation when over-expressed in cells and when synthesized by *in vitro* translation (Levy et al., 2006; Lloyd et al., 2012; Moughamian et al., 2012). Furthermore, dynactin complexes that contain p150/Glued(G59S) exhibit reduced association with dynein in cell lysate (Moughamian et al., 2012). We therefore hypothesized that the enhanced loss-of-function phenotype of the *dnc-1(G33S)* mutant during male meiosis could be due to mis-folded DNC-1(G33S) being unstable and/or unable to interact with dynein. To test these possibilities, we purified three recombinant DNC-1 fragments from bacteria (Fig. 7A). The wild-type control consisted of residues 2-664, which included the CC1 region. The G33S mutation was introduced into this fragment. For DNC-1(ΔN), we purified a corresponding fragment consisting of residues 264-664. The DNC-1 fragments contained an N-terminal 6xHis::MBP tag and a C-terminal StrepTag II that were used in sequential Nickel-NTA and Strep-Tactin affinity purification steps, and the 6xHis::MBP tag was cleaved off with TEV protease after the Nickel-NTA step. Size exclusion chromatography with the purified fragments revealed that the G33S mutant fractionated as the wild-type protein in 150 mM NaCl, displaying no evident aggregation behavior (Fig. 7B). This allowed us to test whether the G33S mutation affects the binding of DNC-1 to dynein intermediate chain (DYCI-1), an interaction that is mediated by p150/Glued’s CC1 in yeast, flies, and vertebrates (McKenney et al., 2011; Morgan et al., 2011; Jie et al., 2015) (Fig. 7C). Using GST pull-down assays with purified recombinant GST-tagged DYCI-1 (residues 1-85), we found that all three DNC-1 fragments bound equally well to the DYCI-1 N-terminus (Fig. 7D). This implies that CAP-Gly domain mis-folding induced by the G33S mutation does not perturb the folding of the neighboring coiled-coil region. To further examine the properties of the purified DNC-1 fragments, we subjected the proteins to high-speed centrifugation (280’000 x g) after a short incubation in different buffers. In 150 mM NaCl, all three fragments remained in the supernatant (Fig. 7E, F). By contrast, decreasing the salt concentration to 100 mM NaCl caused a substantial fraction of the G33S mutant to be recovered in the pellet. A low salt buffer (BRB80) commonly used in microtubule binding assays further decreased the solubility of the G33S mutant, while the wild-type and ΔN fragments remained soluble in all three conditions. We conclude that while both the G33S and ΔN mutant are capable of binding DYCI-1, the G33S mutant is distinct from the ΔN mutant in that it has a strong propensity for aggregation in suboptimal conditions.

**Figure 7.**
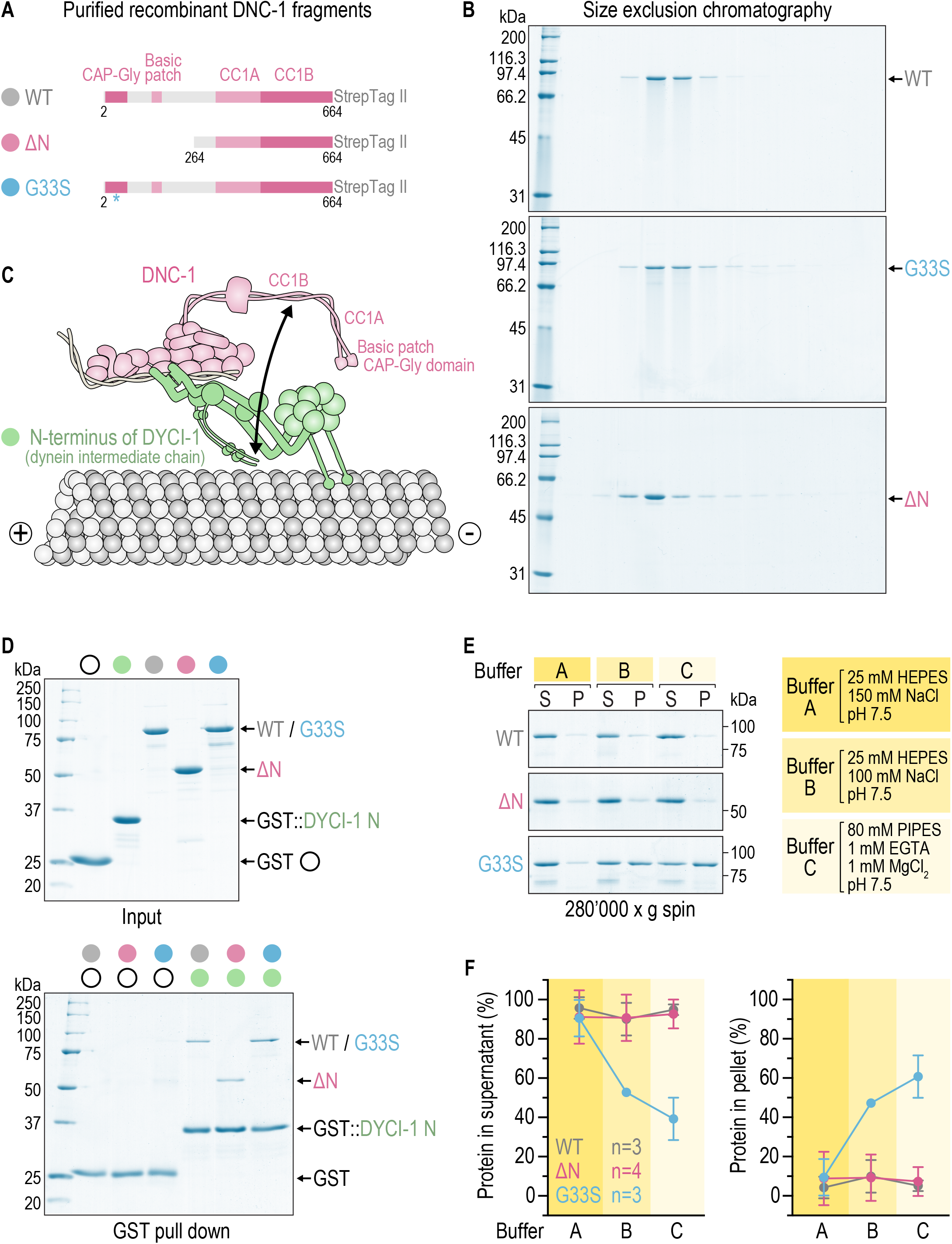
The G33S mutation renders the DNC-1 protein aggregation-prone *in vitro*. **(A)** Schematic of purified recombinant DNC-1 fragments. DNC-1 residue numbers at the fragment N- and C-termini are indicated. **(B)** Coomassie Blue-stained SDS-PAGE gels of purified DNC-1 fragments after size exclusion chromatography. Fractions eluting earlier from the column are to the left. Equivalent elution fractions were loaded for each protein. Molecular weight is in kilodaltons (kDa). **(C)** Cartoon highlighting the interaction between the CC1B region of DNC-1 and the N-terminus of DYCI-1 (dynein intermediate chain), which is examined in (*D*). **(D)** (*top*) Coomassie Blue-stained SDS-PAGE gel of purified proteins used in GST pull down experiments. *DYCI-1 N* denotes residues 1-85. (*bottom*) Coomassie Blue-stained SDS-PAGE gel of proteins eluted from glutathione agarose resin after GST pull-down experiments. Molecular weight is in kilodaltons (kDa). **(E)** Coomassie Blue-stained SDS-PAGE gel of supernatants (S) and pellets (P) after high-speed centrifugation (280’000 x g) of DNC-1 fragments in three different buffers. Buffer C corresponds to the BRB80 buffer commonly used in microtubule binding assays. Molecular weight is in kilodaltons (kDa). **(F)** Fraction of DNC-1 fragments recovered in the supernatant and pellet after high-speed centrifugation, determined by measuring Coomassie Blue intensity on SDS-PAGE gels as in (*E*). *n* denotes the number of independent pelleting experiments.

## DISCUSSION

In this study we took advantage of the *C. elegans* hypomorphic dynactin mutant *dnc-1(G33S)* to address the function of dynein-dynactin in dividing spermatocytes of hermaphrodites and males. The phenotype of *dnc-1(G33S)* shows that dynein-dynactin are involved in prophase centrosome separation and the formation of kinetochore-microtubule attachments. Thus, dynein-dynactin’s roles during spindle assembly in spermatocytes resemble those in the early embryo. In *D. melanogaster*, the only other animal in which dynein function has been inhibited in spermatocytes, mutants of dynein regulators (Asunder, Lis1, NudE) and dynein itself (dynein light chain Tctex) prevent the initial attachment of centrosomes to the prophase nucleus, which results in spindle assembly without centrosomes (Anderson et al., 2009; Wainman et al., 2009; Sitaram et al., 2012). This defect makes it difficult to evaluate dynein’s role at later stages. Because the *dnc-1(G33S)* mutant delays but does not prevent spindle assembly, we were able to define dynactin-dynactin’s contribution to anaphase. A recent analysis of anaphase in *C. elegans* spermatocytes showed that chromosome segregation is accomplished through simultaneous anaphase A and B and that anaphase B accounts for approximately 80% of the total chromosome displacement (Fabig et al., 2020). We find that the *dnc-1(G33S)* mutant essentially eliminates anaphase B, while we occasionally still observed anaphase A-type movement. However, anaphase A by itself is clearly not sufficient for successful chromosome segregation. The essential requirement for dynein-dynactin during male meiotic anaphase contrasts with their relatively minor role during female meiotic anaphase. In oocytes, inhibition of dynein heavy chain, using a fast-acting temperature-sensitive mutant or partial RNAi-mediated depletion, produces defects in spindle morphology and lagging chromosomes in anaphase (Muscat et al., 2015; McNally et al., 2016), but chromosomes that segregate do so with normal kinetics (McNally et al., 2016; Laband et al., 2017). A null mutant of the dynactin p27 subunit (DNC-6) even increases the velocity of chromosome segregation (Laband et al., 2017). The available evidence points to a midzone pushing mechanism for anaphase chromosome movement during female meiosis that is independent of dynein-dynactin and does not involve kinetochores (Dumont et al., 2010; McNally et al., 2016; Laband et al., 2017; Yu et al., 2019). By contrast, the first mitotic division of the *C. elegans* embryo resembles the meiotic division of spermatocytes in that chromosome segregation occurs through anaphase B, which involves dynein-dependent cortical pulling forces. However, chromosomes still segregate successfully in the absence of cortical pulling in the one-cell embryo due to the existence of a normally obscured midzone pushing mechanism (Nahaboo et al., 2015; Yu et al., 2019). Our results with the *dnc-1(G33S)* mutant suggest that such a backup mechanism does not exist in spermatocytes. Indeed, electron tomography analysis by Fabig et al. (2020) failed to detect a typical spindle midzone architecture with overlapping microtubules, and immunofluorescence analysis showed that proteins usually localized to overlapping midzone microtubules in mitotic and female meiotic anaphase do not localize to the midzone region in spermatocytes. The absence of a midzone capable of generating pushing forces predicts that spindle elongation depends exclusively on cortical pulling by dynein-dynactin. Consistent with this, we show that the lack of spindle elongation in the *dnc-1(G33S)* mutant correlates with reduced cortical dynein recruitment. Thus, the reliance on anaphase B and a midzone architecture that differs significantly from that in female meiosis and embryonic mitosis together explain dynein-dynactin’s crucial role in segregating chromosomes in spermatocytes.

Our study highlights the differences between the HMN7B mutation G33S and the two Perry Syndrome mutations (F26L and G45R) that we examined previously (Barbosa et al., 2017): DNC-1 protein levels are higher in Perry Syndrome mutants, tetrapolar spindles in *dnc-1(G45R)* embryos are four fold less frequent than in *dnc-1(G33S)* embryos, and Perry syndrome mutant embryos are mostly viable. In sum, this indicates that DNC-1 with Perry syndrome mutations is more stable than the DNC-1(G33S) protein, which is consistent with prior work on the human and *D. melanogaster* orthologs (Ahmed et al., 2010; Lloyd et al., 2012; Moughamian et al., 2012). A notable aspect of the *dnc-1(G33S)* mutant is that the phenotypic severity varies significantly between dividing spermatocytes and the dividing early embryo (discounting the tetrapolar spindle phenotype in the one-cell embryo, which is a consequence of defects in spermatogenesis). This is not because dynactin is less important for mitosis, since centrosome separation, and therefore chromosome segregation, is completely inhibited in the early embryo when DNC-1 is depleted by RNAi. Nor is it the case that DNC-1’s CAP-Gly domain has a more important function in spermatocytes than the early embryo, since removal of the CAP-Gly domain in the *dnc-1(ΔN)* mutant only mildly impacts cell division in either tissue. Although it was impracticable to directly compare DNC-1 protein levels in the two tissues, quantitative analysis of the mutant phenotypes implies that DNC-1(G33S) levels are lower than DNC-1(ΔN) levels in dividing spermatocytes, while mutant protein levels are similar in the dividing early embryo. The idea that DNC-1 (G33S) is inherently unstable is supported by characterization of the mutant protein *in vitro*. We show that a recombinant DNC-1(G33S) fragment that includes the CC1 region can be purified in a nonaggregated state, in contrast to the human p150/Glued CAP-Gly domain fragment containing the G59S mutation, which was reported to be insoluble when expressed in bacteria (Ahmed et al., 2010). Efficient binding of the G33S fragment to the N-terminus of dynein intermediate chain suggests that the point mutation does not directly interfere with the interaction between dynactin and dynein *in vivo*. However, the G33S fragment readily aggregates in conditions that have no effect on the solubility of the wild-type and ΔN fragments. We speculate that spermatocytes and the early embryo differ in aspects that determine DNC-1(G33S) solubility, such as in the content and concentration of molecular chaperones or in post-translational modifications on DNC-1 that may modulate aggregation behavior. The *dnc-1(G33S)* mutant illustrates that for mutations that induce mis-folding in an ubiquitously expressed protein, the extent to which protein function is lost can vary significantly between cell types because the mutant protein may be more stable in some cellular environments than others. Whether such an effect could possibly contribute to the neuronal cell-type specificity of HMN7B remains an interesting open question.

## MATERIALS AND METHODS

### *C. elegans* strains

Worm strains (Table S1) were maintained at 20 °C on standard NGM plates seeded with OP50 bacteria. The *dnc-1(ΔN)* mutant and the *FL* and *Δexon4* variants of *dnc-1(G33S)* were generated by CRISPR/Cas9-mediated genome editing, as described previously (Arribere et al., 2014; Paix et al., 2014; Barbosa et al., 2017). Genomic sequences targeted by sgRNAs are listed in Table S2. The modifications were confirmed by sequencing and strains were outcrossed 6 times against the wild-type N2 strain. Other fluorescent markers were subsequently introduced by mating. The *dnc-1(G33S)* mutant was maintained in a heterozygous state using the GFP-marked balancer nT1 [qIs51]. Homozygous F1 progeny from balanced heterozygous mothers were identified by the absence of GFP fluorescence. Males of the *dnc-1(G33S)* mutant were generated by heat-shocking heterozygous L4 hermaphrodites for 6 h at 30°C. Males were maintained by mating 30 heterozygous males with 15 heterozygous hermaphrodites for 24 h at 20 °C, and homozygous male progeny was identified four days later by the absence of GFP fluorescence.

### Embryonic viability and egg laying assay

Embryonic viability tests were performed at 20 °C. L4 hermaphrodites were grown on NGM plates with bacteria for 40 h at 20 °C and singled-out to NGM plates with a small amount of bacteria. After 8 h, mothers were removed and the number of hatched and unhatched embryos on each plate was determined 16 h later.

### Life span assay

Animals hatched from eggs laid by mothers in a 2-h window (day 0) were grown at 20 °C and transferred every 2 days to a new NGM plate with bacteria. Animals were scored as alive or dead every 1 - 3 days. Animals were considered dead if they did not respond when touched with a platinum wire and if there was no evidence of pharyngeal pumping. Animals that were found dead on the edge of the plate or escaped were excluded from the assay.

### Reverse transcription PCR

Total RNA was isolated from adult hermaphrodites using the TRIzol Plus RNA Purification Kit (Invitrogen). After 3 washes with M9 buffer (86 mM NaCl, 42 mM Na_2_HPO_4_, 22 mM KH_2_PO_4_, 1 mM MgSO_4_), pelleted worms were homogenized in 200 μL of TRIzol reagent with a pellet pestle homogenizer and incubated at room temperature for 5 min. After addition of 40 μL chloroform, samples were shaken vigorously by hand, incubated at room temperature for 3 min, and centrifuged at 12’000 x g for 15 min at 4 °C. The upper phase containing the RNA was transferred to an RNase-free tube and an equal volume of 70 % ethanol was added. Further RNA purification steps were performed according to the manufacturer’s instructions. Purified RNA was treated with DNase I (Thermo Scientific), and cDNA was synthesized with the iScript Select cDNA Synthesis Kit (Bio-Rad Laboratories). The following oligos were used for the PCR reactions: forward oligo on *dnc-1* exon 3 (GAATGTCACCTGCTGCTT); forward oligo on *dnc-1* exon 4 (AAAGCGGTCTACAACTCC); forward oligo on *dnc-1* exon 7 (ATGACCGAGTCCGGCACAGA); reverse oligo on *dnc-1* exon 5 (GATTGCGATAAGTTGGAGA); reverse oligo on *dnc-1* exon 6 (AGTAGTCGTGGACGCTTT); reverse oligo on *dnc-1* exon 8 (CTCGTCTTGCCATGCTTGTA); forward oligo on *gpd-2* exon 1-2 (TCAAGGTCTACAACTCAAG); reverse oligo on *gpd-2* exon 3 (GATGGAGCAGAGATGATG).

### RNA interference

dsRNA against *dnc-1* (Gama et al., 2017) was delivered to L4 hermaphrodites by injection. Injected animals were kept on NGM plates seeded with bacteria for 48 h at 20 °C before embryos were isolated for live imaging.

### Immunoblotting

200 adult hermaphrodites, or L3 hermaphrodites for the *dnc-1(null)* mutant, were collected into 1 mL of M9 buffer, washed with 3 x 1 mL M9 buffer, and once with M9 buffer containing 0.05 % Triton X-100. To 100 μL of worm suspension, 33 μL 4x SDS PAGE sample buffer [250 mM Tris-HCl pH 6.8, 30 % (v/v) glycerol, 8 % (w/v) SDS, 200 mM DTT, 0.04 % (w/v) bromophenol blue] and 20 μL of glass beads were added. Samples were incubated for 3 min at 95 °C and vortexed for 10 min with intermittent heating. After centrifugation at 20’000 x g for 1 min at room temperature, the protein fraction in the supernatant was resolved on a 10 % SDS-PAGE gel and transferred to a 0.2-μm nitrocellulose membrane (Amersham, GE Healthcare). The membrane was blocked with 5 % (w/v) non-fat dry milk in TBST (20 mM Tris, 140 mM NaCl, 0.1 % Tween, pH 7.6) and incubated overnight at 4 °C in TBST with rabbit anti-DNC-1 antibody GC2 (1:1’000; Gama et al., 2017), rabbit anti-DNC-2 antibody GC5 (1:5’000; Gama et al., 2017), rabbit anti-DYCI-1 antibody (1:1’000; Barbosa et al., 2017), or mouse anti-α-tubulin antibody B512 (1:5’000; Sigma-Aldrich). The membrane was rinsed 5 x with TBST and incubated in TBST with goat secondary antibodies coupled to HRP (1:10’000; Jackson ImmunoResearch) for 1 h at room temperature. After 3 washes with TBST, proteins were visualized by chemiluminescence using Pierce ECL Western Blotting Substrate (Thermo Fisher Scientific) and x-ray film (Amersham, GE Healthcare).

### Immunofluorescence

10 - 12 gravid adult hermaphrodites were dissected into 3 μL of M9 buffer on a poly-*L*-lysine-coated slide by performing a scissor motion with two 25-gouge needles. A 13-mm^2^ round coverslip was placed on the drop, and slides were plunged into liquid nitrogen. After rapid removal of the coverslip (“freeze-cracking”), samples were fixed for 20 min in −20 °C methanol. Samples were re-hydrated for 2 x 5 min in PBS (137 mM NaCl, 2.7 mM KCl, 8.1 mM Na2HPO4, 1.47 mM KH2PO4), blocked with AbDil (PBS, 2 % BSA, 0.1 % Triton X-100) in a humid chamber for 30 min at room temperature, and incubated in AbDil with rabbit anti-SAS-4 antibody (1:1’000; Dammermann et al., 2004) overnight at 4 °C. After washing for 4 x 5 min in PBS, samples were incubated in AbDil with Alexa Fluor 594 goat anti-rabbit IgG (1:300; Jackson ImmunoResearch) for 1 h at room temperature. Samples were washed for 4 x 5 min in PBS and mounted in Prolong Gold with DAPI stain (Invitrogen). Imaging was performed on an Axio Observer microscope (Zeiss) equipped with an Orca Flash 4.0 camera (Hamamatsu) and an HXP 200C Illuminator (Zeiss), controlled by ZEN 2.3 software (Zeiss). Images were captured with a 100x NA 1.46 Plan-Apochromat objective at 1 x 1 binning.

### Live imaging

#### Locomotion assay

Animals synchronized at the L1 stage were grown at 20 °C and transferred every 2 days to a new NGM plate with bacteria. The 72-h time point after release from L1 arrest was considered adult day 1. For imaging, animals were transferred to a slide containing a 2-μL drop of M9 buffer. Movements were tracked at 20 °C for 1 min at 40 frames per second using a SMZ 745T stereoscope (Nikon) equipped with a QIClic CCD camera (QImaging) and controlled by Micro-Manager software (Open Imaging). The wrMTrck plugin for Image J was used for automated counting of body bends.

#### Differential interference contrast (DIC) imaging of the one-cell embryo

Gravid adult hermaphrodites were dissected in a watch glass filled with 0.7x Egg Salts medium (1x Egg Salts is 118 mM NaCl, 40 mM KCl, 3.4 mM MgCl2, 3.4 mM CaCl2, 5 mM HEPES, pH 7.4). Embryos were mounted onto a fresh 2 % agarose pad and covered with an 18 mm x 18 mm coverslip (No. 1.5H, Marienfeld). Imaging was performed at 20 °C on the Axio Observer microscope described above with a 63x NA 1.4 Plan-Apochromat objective at 2 x 2 binning.

#### Fluorescence imaging of the early embryo

Embryos co-expressing mCherry::HIS-11 and GFP::TBB-2 or expressing GFP::DHC-1 were isolated and mounted as described above. Imaging was performed at 20 °C on a Nikon Eclipse Ti microscope coupled to an Andor Revolution XD spinning disk confocal system, composed of an iXon Ultra 897 CCD camera (Andor Technology), a solid-state laser combiner (ALC-UVP 350i, Andor Technology), and a CSU-X1 confocal scanner (Yokogawa Electric Corporation). The system was controlled by Andor IQ3 software (Andor Technology). Images were acquired with a 60x NA 1.4 Plan-Apochromat objective at 1 x 1 binning.

#### Fluorescence imaging of dividing spermatocytes

L4 hermaphrodites or young adult males were paralyzed with in a 5-μL drop of 5 mM levamisole in M9 buffer for 10 min on an 18 mm x 18 mm coverslip and mounted on a fresh 5 % agarose pad. Imaging was performed on the spinning disk confocal system described above with a 60x NA 1.4 Plan-Apochromat objective at 1 x 1 binning.

### Image acquisition and analysis

#### Migration of the male pronucleus

DIC time-lapse sequences, consisting of a 3 x 1.5 μm z-stack captured every 5 s, were recorded from the start of pronuclear migration until the onset of cytokinesis. Embryo length was defined as the distance between the outermost points of the egg shell with the anterior reference point set to 0 %. Using the un-projected images, the XY coordinates of the male pronucleus were determined in each frame using the MTrackJ plugin of the Fiji software (Image J version 2.0.0-rc-56/1.52p) by manually clicking in the center of the nucleus. The relative position of the pronucleus along the anterior-posterior axis was calculated, and tracks from individual embryos were aligned relative to nuclear envelope breakdown or pronuclear meeting.

#### Frequency of tetrapolar spindle formation in the one-cell embryo

DIC time-lapse sequences of the first embryonic division, acquired as described above, were manually inspected to determine the number of spindle poles.

#### Cell division in the multicellular embryo

Fluorescence time-lapse sequences (mCherry::HIS-11/GFP::TBB-2), consisting of a 11 x 1 μm z-stack captured every 30 s, were recorded for 1 h starting just prior to the onset of anaphase in the EMS cell of the four-cell embryo.

#### Pole-pole distance in the one-cell embryo

Fluorescence time-lapse sequences (mCherry::HIS-11/GFP::TBB-2), consisting of a 11 x 1 μm z-stack captured every 20 s, were recorded from the start of pronuclear migration until the onset of cytokinesis. The XYZ coordinates of each centrosome were determined in every frame using Imaris software (Oxford instruments). All tracks automatically generated by Imaris were manually inspected. The pole-pole distance was calculated in 3D, and tracks from individual embryos were aligned relative to anaphase onset.

#### Pole-pole distance in meiosis I spermatocytes

Fluorescence time-lapse sequences (mCherry::HIS-11/GFP::TBB-2), consisting of a 35 x 0.33 μm z-stack captured every 30 s, were recorded in the meiotic division zone during 1 h. The XYZ coordinates of centrosomes were determined in every frame for selected spermatocytes using Imaris. All tracks automatically generated by Imaris were manually inspected. The pole-pole distance was calculated in 3D, and tracks from individual embryos were aligned relative to pole splitting in late anaphase I.

#### GFP::DHC-1 signal in the one-cell embryo

Fluorescence time-lapse sequences, consisting of a 8 x 1 μm z-stack captured every 20 s, were recorded from 40-50 s prior to pronuclear meeting until the onset of cytokinesis. Nuclear envelope (NE) signal was quantified 3 frames prior to pronuclear meeting using a maximum intensity projection of the 3 z-sections representing the best in-focus images of the NE. A 2 pixel-wide line was drawn on top of the NE along its entire circumference, and a similar line was drawn next to the NE on the cytoplasmic side around the nucleus. The mean fluorescence signal of the cytoplasmic line was subtracted from the mean fluorescence signal of the NE line.

Kinetochore signal was measured 7-8 frames before the onset of sister chromatid separation using a maximum intensity projection of the z-stack. The top 10 local maxima intensities on kinetochores were identified using the “Find Maxima” function in Fiji, and the 10 values were averaged. The mean fluorescence intensity of the spindle background close to the kinetochore region was measured and subtracted from the kinetochore signal.

Mitotic spindle signal was measured 2 frames after the onset of chromosome segregation using a maximum intensity projection of the z-stack. The mean intensity in 3 separate 10 x 10 pixel squares on the spindle was determined and the three values were averaged. The mean intensity of three equivalent squares in the cytoplasm adjacent to the spindle was subtracted from the spindle signal.

#### GFP::DHC-1 signal at the EMS-P2 cell boundary in the four-cell embryo

Fluorescence time-lapse sequences, consisting of a 11 x 1 μm z-stack captured every 30 s, were recorded from the beginning of nuclear envelope breakdown in AB cells until cytokinesis of the P2 cell. The signal at the EMS-P2 cell boundary was measured at the time of EMS spindle rotation in maximum intensity projections of the z-stacks. The top 10 local maxima intensities on the EMS-P2 cell boundary were identified using the “Find Maxima” function in Fiji, and the 10 values were averaged. The mean fluorescence intensity of an adjacent area (20 x 20 pixels) in the EMS cytoplasm was subtracted from the signal at the EMS-P2 cell boundary.

#### Cortical GFP::DHC-1 signal in spermatocytes

Single 15 x 0.5 μm z-stacks were captured in the meiotic division zone. Cortical signal was quantified in a single z-section representing the best in-focus image of cell-cell boundaries. A 2 pixel-wide line was drawn on top of the cell-cell boundary, and a similar line was drawn next to the cell-cell boundary on the cytoplasmic side of adjacent cells. The mean fluorescence signal of the cytoplasmic line was subtracted from the mean fluorescence signal of the cell-cell boundary line.

#### GFP::DHC-1 in dividing spermatocytes

Fluorescence time-lapse sequences, consisting of a 15 x 0.5 μm z-stack captured every 30 s, were recorded during 1 h in the meiotic division zone.

### Constructs for biochemistry

The cDNA of *dnc-1* corresponding to residues 2-664 (wild type and G33S) or 264-664 (ΔN) was cloned into a 2CT expression vector containing N-terminal 6xHis::maltose-binding protein (MBP) followed by a linker with a TEV protease cleavage site and a C-terminal linker followed by the Strep-tag II. The cDNA of DYCI-1 (residues 1-85) was cloned into pGEX-6P-1 containing N-terminal GST and a C-terminal flexible linker followed by 6xHis.

### Protein expression and purification

Expression vectors were transformed into the *E. coli* strain BL21. Expression was induced at an OD_600_ of 0.9 with 0.1 mM IPTG. After expression overnight at 18 °C, cells were harvested by centrifugation at 4’000 x g for 20 min.

For purification of DNC-1::Strep-tag II proteins, bacterial pellets were resuspended in lysis buffer B (50 mM HEPES, 250 mM NaCl, 10 mM imidazole, 0.1 % Tween 20, 1 mM DTT, 1 mM PMSF, 2 mM benzamidine-HCl, 1 mg/mL lysozyme, pH 8.0), disrupted by sonication, and cleared by centrifugation at 34’000 x g for 40 min. Proteins were purified by tandem affinity chromatography using HisPur Ni-NTA resin (Thermo Fisher Scientific) followed by Strep-Tactin Sepharose resin (IBA). Ni-NTA resin was incubated in batch with the cleared lysate and washed with wash buffer A (25 mM HEPES, 250 mM NaCl, 25 mM imidazole, 0.1 % Tween 20, 1 mM DTT, 2 mM benzamidine-HCl, pH 8.0). Proteins were eluted on a gravity column with elution buffer B (50 mM HEPES, 150 mM NaCl, 250 mM imidazole, 1 mM DTT, 2 mM benzamidine-HCl, pH 8.0). Fractions containing the recombinant proteins were pooled, incubated overnight with TEV protease to cleave off 6xHis::MBP, incubated in batch with Strep-Tactin Sepharose resin, and washed with wash buffer B (25 mM HEPES, 250 mM NaCl, 0.1 % Tween 20, 1 mM DTT, 2 mM benzamidine-HCl, pH 8.0). Proteins were eluted on a gravity column with elution buffer E (100 mM Tris-HCl, 150 mM NaCl, 1 mM EDTA, 2.5 mM desthiobiotin, pH 8.0; IBA). Fractions containing the recombinant proteins were pooled and dialyzed against storage buffer (25 mM HEPES, 150 mM NaCl, pH 7.5). Glycerol and DTT were added to final concentrations of 10 % (v/v) and 1 mM, respectively, and aliquots were flash frozen in liquid nitrogen and stored at −80°C.

For purification of GST::DYCI-1(1-85)::6xHis and GST::6xHis (negative control for the pull downs), bacterial pellets were resuspended in lysis buffer A (50 mM HEPES, 250 mM NaCl, 0.1 % Tween 20, 10 mM EDTA, 10 mM EGTA, 1 mM DTT, 1 mM PMSF, 2 mM benzamidine-HCl, 1 mg/mL lysozyme, pH 8.0), disrupted by sonication, and cleared by centrifugation at 34’000 x g for 40 min. Proteins were purified by tandem affinity chromatography using Glutathione Agarose resin (Thermo Fisher Scientific) followed by Ni-NTA resin. Glutathione Agarose resin was incubated in batch with the cleared lysate, washed with wash buffer B (25 mM HEPES, 250 mM NaCl, 0.1 % Tween 20, 1 mM DTT, 2 mM benzamidine-HCl, pH 8.0), and proteins were eluted on a gravity column with elution buffer A (50 mM HEPES, 150 mM NaCl, 10 mM reduced L-glutathione, 1 mM DTT, 2 mM benzamidine-HCl, pH 8.0). Fractions containing the recombinant proteins were pooled, incubated in batch with Ni-NTA resin, washed with wash buffer A (25 mM HEPES, 250 mM NaCl, 25 mM imidazole, 0.1% Tween 20, 1 mM DTT, 2 mM benzamidine-HCl, pH 8.0), and proteins were eluted on a gravity column with elution buffer B (50 mM HEPES, 150 mM NaCl, 250 mM imidazole, 1 mM DTT, 2 mM benzamidine-HCl, pH 8.0). GST::6His and GST::DYCI-1(1-85)::6xHis were further purified by size-exclusion chromatography on a Superose 6 10/300 column equilibrated with storage buffer (25 mM HEPES, 150 mM NaCl, pH 7.5). Glycerol and DTT were added to final concentrations of 10 % (v/v) and 1 mM, respectively, and aliquots were flash frozen in liquid nitrogen and stored at −80°C.

### Size exclusion chromatography

80 μg of the purified recombinant DNC-1 fragments were loaded on a Superose 6 10/300 column (GE Healthcare) equilibrated with storage buffer (25 mM HEPES, 150 mM NaCl, pH 7.5). Size exclusion chromatography was performed in storage buffer at room temperature on an ÄKTA Pure 25L1 system (GE Healthcare). Elution of proteins was monitored at 280 nm. 20 μL of successive 0.5-mL elution fractions were separated on a 10 % SDS-PAGE gel and proteins were visualized by Coomassie Blue staining.

### GST pull-down assay

100 pmol of the purified recombinant DNC-1 fragments were incubated for 1 h at 4 °C with 50 pmol of purified recombinant GST::DYCI-1(1-85)::6xHis or GST::6xHis in 150 μL pull-down buffer (50 mM HEPES, 100 mM NaCl, 0.1 % Tween 20, 5 mM DTT, pH 7.5) containing 15 μL of Glutathione Agarose resin. The resin was washed with 3 x 500 μL of pull-down buffer, and proteins were eluted with pull-down buffer containing 15 mM reduced L-glutathione for 15 min at room temperature. Eluted proteins were separated on a 10 % SDS-PAGE gel and visualized by Coomassie Blue staining.

### Sedimentation assay

50 pmol of the purified recombinant DNC-1 fragments were added to buffer A (25 mM HEPES, 150 mM NaCl, pH 7.5), buffer B (25 mM HEPES, 100 mM NaCl, pH 7.5), or buffer C (80 mM PIPES, 1 mM EGTA, 1 mM MgCl2, pH 7.5) and incubated for 10 min at 25 °C. Samples were pelleted at 280’000 x g for 10 min at 25 °C in an Optima XP centrifuge with a TLA-100 rotor (Beckman-Coulter). Supernatant and pellet fractions were separated on a 10 % SDS-PAGE gel and proteins were visualized by Coomassie Blue staining. Protein band intensities were quantified in Fiji by drawing a box around the protein band and two same-sized boxes above and below. The integrated signal of the two background boxes was averaged and subtracted from the integrated signal of the protein box.

### Statistical analysis

Statistical analysis was performed with Prism 8.0 software (GraphPad). Values in figures and text are reported as mean with a 95 % confidence interval of the SEM (mean ± 95 % CI). After normality tests, data was analyzed by one-way ANOVA followed by Dunnett’s multiple comparison test or by ANOVA on ranks (Kruskal-Wallis nonparametric test) followed by Dunn’s multiple comparison test, as indicated in the figure legends.

## ACKNOWLEDGEMENTS

The authors thank Alex Dammermann for the SAS-4 antibody. This study was financed by the Fundação para a Ciência e a Tecnologia (FCT)/Ministério da Ciência, Tecnologia e Ensino Superior through project grant PTDC/BIA-CEL/30507/2017. R. G. and A. X. C. have FCT Principal Investigator positions (CEECIND/00333/2017 and CEECIND/01967/2017, respectively). D. J. B. has an FCT Junior Researcher position (DL 57/2016/CP1355/CT0007).

---

**Figure S1.**
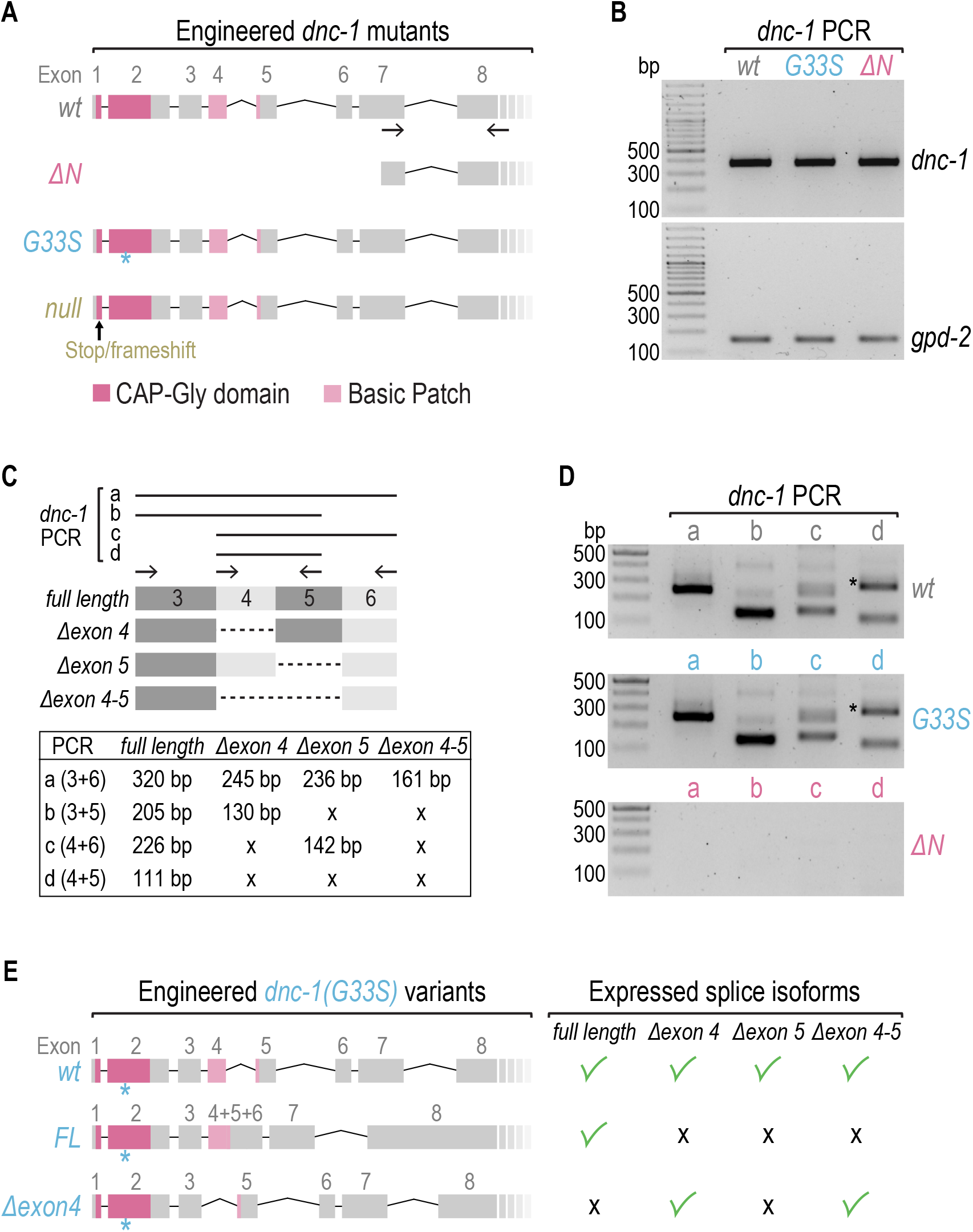
mRNA levels and splice isoforms of *dnc-1* mutants. **(A)** Schematic of the genomic *dnc-1* locus before (*wt*) and after CRISPR/Cas9-mediated editing. The location of the primer pair used to assess mRNA levels in (*B*) is indicated below the *wt* sequence. **(B)** DNA gel of reverse transcription PCR products. The template for the reverse transcription reaction was total RNA isolated from adult hermaphrodites. *gpd-2* (GAPDH) serves as the loading control. DNA size marker is a 100-bp ladder. **(C)** Primer pairs for the detection of *dnc-1* splice isoforms used for the reverse transcription PCRs shown in (*D*). Crosses (x) indicate that the PCR will not amplify any product. **(D)** DNA gels of reverse transcription PCR products. The asterisk denotes an unspecific PCR product of primer pair “d”. DNA size marker is a 100-bp ladder. **(E)** Schematic of the genomic *dnc-1* locus after CRISPR/Cas9-mediated editing to generate specific *dnc-1* splice variants in the context of the G33S mutation. Exon fusion or deletion restricted *dnc-1* expression to the full-length isoform (*FL*) or to isoforms that lack exon 4 *(Δexon4)*, respectively.

**Figure S2.**
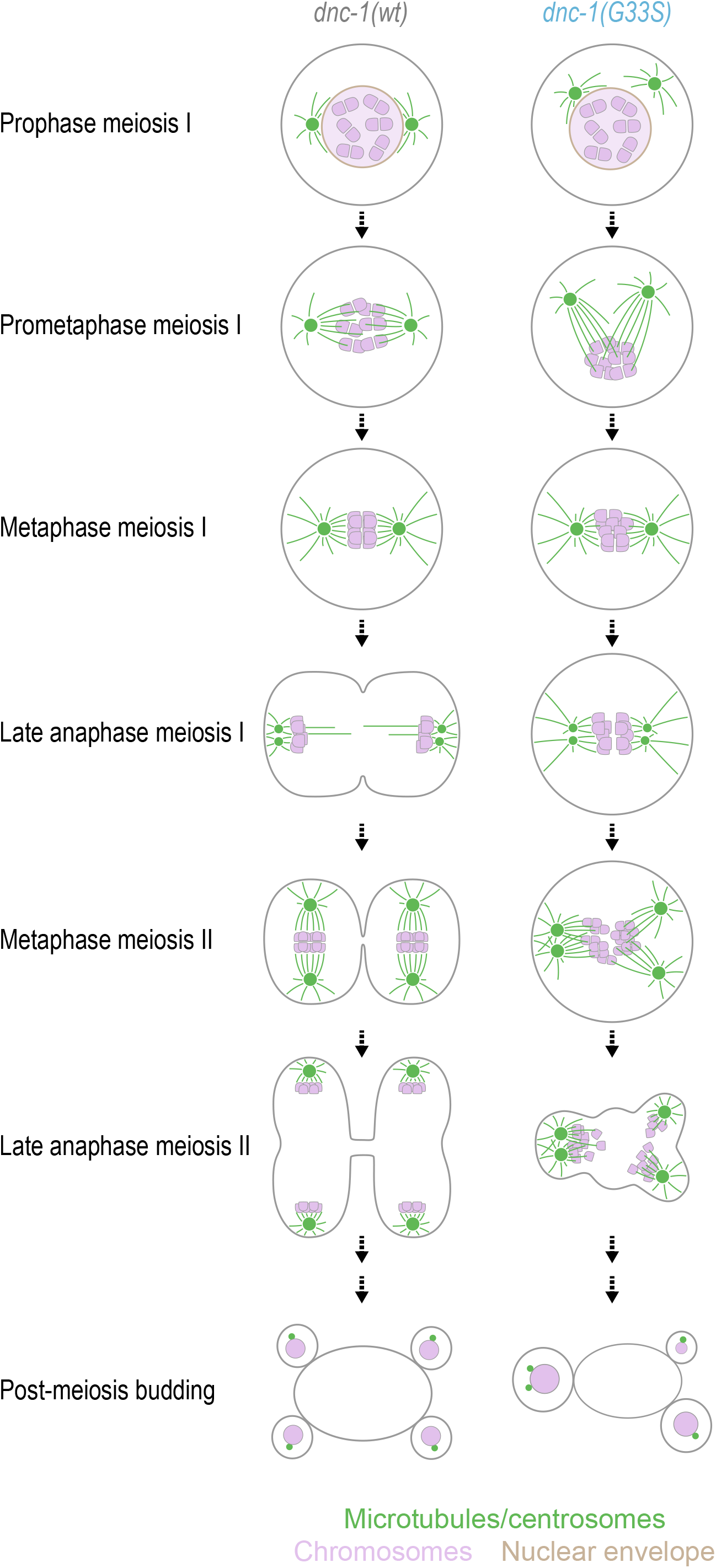
Cartoon summary of defects in *dnc-1(G33S)* spermatocytes. In the *dnc-1(G33S)* mutant, spermatocytes essentially skip meiosis I and exhibit a highly aberrant meiosis II, which results in aneuploid sperm that contains one or two centrosomes.

**Figure S3.**
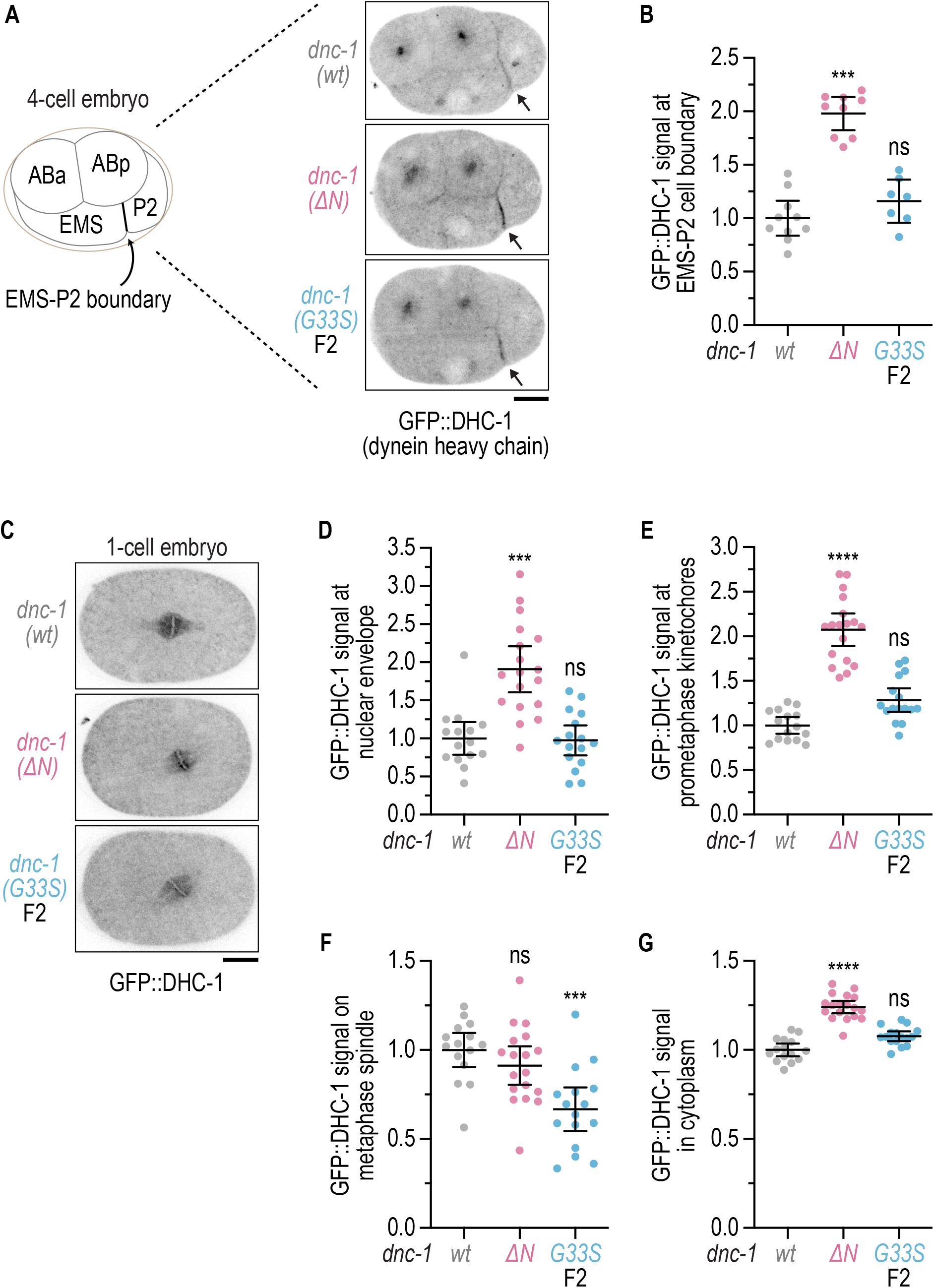
Subcellular dynein localization is only minimally affected in *dnc-1(G33S)* embryos. **(A)** Fluorescence images of GFP::DHC-1 in the four-cell embryo. Cortical dynein is significantly enriched at the EMS-P2 boundary (arrow). Scale bar, 10 μm. **(B)** Cortical GFP::DHC-1 signal (mean ± 95 % CI) measured at the EMS-P2 boundary, as shown in *(A)*, and normalized to the wild-type control (*wt*). Statistical significance (wild-type versus mutant *dnc-1)* was determined by ANOVA on ranks (Kruskal-Wallis nonparametric test) followed by Dunn’s multiple comparison test. \*\*\**P* < 0.001; ns = not significant, *P* > 0.05. **(C)** Fluorescence images of GFP::DHC-1 in the metaphase one-cell embryo. Scale bar, 10 μm. **(D) - (G)** GFP::DHC-1 signal (mean ± 95 % CI) measured at the nuclear envelope (*D*), on prometaphase kinetochores (*E*), on the metaphase spindle (*F*), and in the cytoplasm (*G*), normalized to the wild-type control (*wt*). Statistical significance (wild-type versus mutant *dnc-1*) was determined by ANOVA on ranks (Kruskal-Wallis nonparametric test) followed by Dunn’s multiple comparison test. \*\*\*\**P* < 0.0001; \*\*\**P* < 0.001; ns = not significant, *P* > 0.05.

**Table S1:** *C. elegans* strains.

| Strain | Genotype |
| --- | --- |
| N2 | Wild type (ancestral N2 Bristol) |
| GCP231 | *dnc-1*(prt12 [G33S]) IV/nT1 [qls51] (IV;V) |
| GCP245 | *dnc-1*(prt18 [G45R]) IV |
| GCP255 | *dnc-1*(prt23 [null]) IV/nT1 [qls51] (IV;V) |
| GCP304 | *dnc-1*(prt32 [Δaa1-263]) IV |
| GCP675 | *dnc-1*(prt32 [Δaa1-263]) IV; *ijmSi8* [pJD362; *pmex-5::egfp::tbb-2*; *mcherry::his-11*; *cb-unc-119(+)*] II; *unc-119*(ed3) III?* |
| GCP706 | *dnc-1*(prt12 [G33S]) IV/nT1 [qls51] (IV;V); *ijmSi8* [pJD362; *pmex-5::egfp::tbb-2*; *mcherry::his-11*; *cb-unc-119(+)*] II; *unc-119*(ed3) III?* |
| GCP726 | *dnc-1*(prt115 [G33S]) *dnc-1*(prt8 [exon 5-6 fusion]) *dnc-1*(prt20 [exon 4-5 fusion]) IV/nT1 [qls51] (IV;V) |
| GCP771 | *dnc-1*(prt116 [G33S]) *dnc-1*(prt9 [Δexon4]) IV/nT1 [qls51] (IV;V) |
| GCP853 | *dnc-1*(prt32 [Δaa1-263]) IV; *dhc-1*(he264 [N-terminal egfp]) I |
| GCP891 | *dnc-1*(prt12 [G33S]) IV/nT1 [qls51] (IV;V); *dhc-1*(he264 [N-terminal egfp]) I |
| JDU21 | *ijmSi8* [pJD362; *pmex-5::egfp::tbb-2*; *mcherry::his-11*; *cb-unc-119(+)*] II; *unc-119*(ed3) III?* |
| SV1803 | *dhc-1*(he264 [N-terminal egfp]) I |

\* *unc-119*(ed3) III? was present in the paternal strain, but this strain has not been sequenced to determine if the ed3 mutation is present in the *unc-119* gene.

**Table S2:**
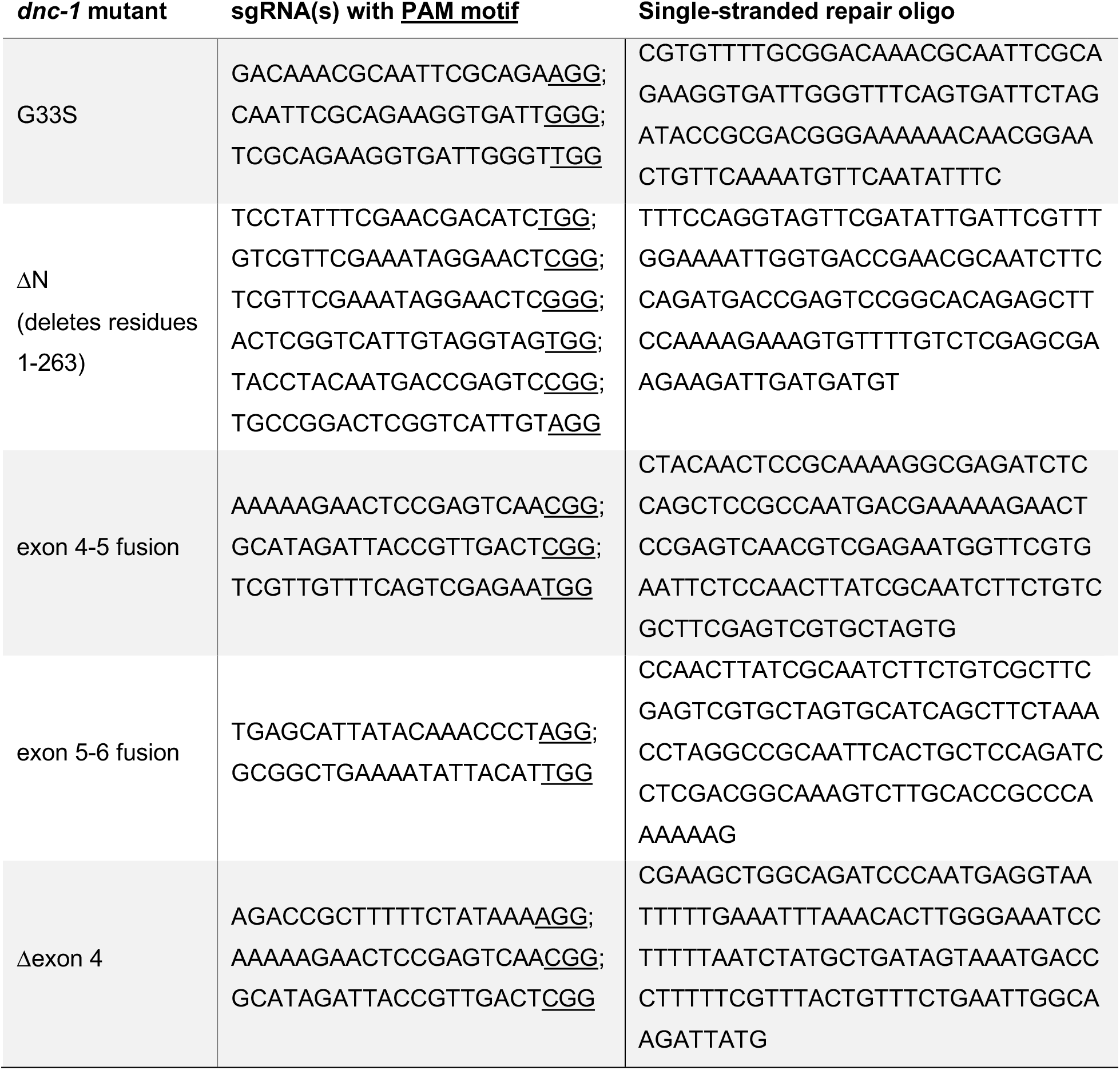
sgRNAs and repair templates for CRISPR/Cas9-mediated genome editing.

